# Simulation of adaptive immune receptors and repertoires with complex immune information to guide the development and benchmarking of AIRR machine learning

**DOI:** 10.1101/2023.10.20.562936

**Authors:** Maria Chernigovskaya, Milena Pavlović, Chakravarthi Kanduri, Sofie Gielis, Philippe A. Robert, Lonneke Scheffer, Andrei Slabodkin, Ingrid Hobæk Haff, Pieter Meysman, Gur Yaari, Geir Kjetil Sandve, Victor Greiff

## Abstract

Machine learning (ML) has shown great potential in the adaptive immune receptor repertoire (AIRR) field. However, there is a lack of large-scale ground-truth experimental AIRR data suitable for AIRR-ML-based disease diagnostics and therapeutics discovery. Simulated ground-truth AIRR data are required to complement the development and benchmarking of robust and interpretable AIRR-ML methods where experimental data is currently inaccessible or insufficient. The challenge for simulated data to be useful is incorporating key features observed in experimental repertoires. These features, such as antigen or disease-associated immune information, cause AIRR-ML problems to be challenging. Here, we introduce LIgO, a software suite, which simulates AIRR data for the development and benchmarking of AIRR-ML methods. LIgO incorporates different types of immune information both on the receptor and the repertoire level and preserves native-like generation probability distribution. Additionally, LIgO assists users in determining the computational feasibility of their simulations. We show two examples where LIgO supports the development and validation of AIRR-ML methods: (1) how individuals carrying out-of-distribution immune information impacts receptor-level prediction performance and (2) how immune information co-occurring in the same AIRs impacts the performance of conventional receptor-level encoding and repertoire-level classification approaches. LIgO guides the advancement and assessment of interpretable AIRR-ML methods.

## Introduction

B- and T-cell receptors (BCRs and TCRs, or AIRs — adaptive immune receptors) are the agents of the adaptive immune system that recognize and neutralize antigens, such as bacteria, viruses, cancer cells, or autoantigens. Immune memory of past and ongoing immune states, i.e. different disease or infection states, is stored in AIR repertoires (AIRRs). In the manuscript text, we will interchangeably use the terms AIRR and repertoire, as well as receptor and AIR. The immune memory is stored as immune information or “immune signal”, which may be encoded in AIRR frequency, e.g., clonal expansion (Greiff et al. 2015), receptor sequence, e.g., antigen-specific binding motifs or disease-associated motifs where motifs may be entire sequences (Emerson et al. 2017) or subsequences (Akbar et al. 2021; Ostmeyer et al. 2019; Shrock et al. 2023) or as a combination thereof. The complexity of AIRR biology contributes to the long-standing problem of how to extract and use these immune signals to understand the mechanisms of adaptive immunity and utilize them for immunodiagnostics (Arnaout et al. 2021; Emerson et al. 2017; Vujkovic et al. 2023) and immunotherapeutic design (Greiff, Yaari, and Cowell 2020; Akbar, Bashour, et al. 2022).

The recognition of antigens can be defined at the receptor level and at the repertoire level. On the receptor level, each immune receptor is specific to one or more antigens, and is potentially polyreactive to several antigens or pathogens using several possible paratopes (Wucherpfennig et al. 2007; Robert, Marschall, and Meyer-Hermann 2018; Garrett Rappazzo et al. 2023). This makes antigen recognition difficult to quantitatively describe because each antigen can be bound by many receptors with very little sequence overlap (Robert, Marschall, and Meyer-Hermann 2018; Boughter et al. 2020; Mason et al. 2021; Glanville et al. 2017; Dash et al. 2017; Straub et al. 2023; Dorigatti et al. 2023). On the repertoire level, immune recognition is determined by a set of antigen-specific AIRs and usually, it remains unknown which AIRs from the whole repertoire are specific to the target and how the distribution (e.g., frequency, positional weight matrices) of antigen-specific AIRs changes across immune states. Recognition on the repertoire level is also complex to describe and quantify because each individual possesses a different set of AIRs with a genetic sequence diversity of about 10^8^–10^9^ different AIRs per individual and low sequence overlap (<1%) across individuals (Greiff, Yaari, and Cowell 2020; Greiff, Menzel, et al. 2017; Greiff, Weber, et al. 2017). Altogether, repertoire-level research problems range from the identification of repertoires of a given immune state containing immune-state-associated AIR biomarkers to the identification of the specific receptors within a repertoire that confers an immune recognition property to this repertoire. Furthermore, as few as one AIR per million to several thousands may be associated with a disease in naive and non-naive repertoires (Christophersen et al. 2014; Abbott et al. 2018), indicating that the immune signal may be very sparingly represented (Brown et al. 2019; C. R. Weber et al. 2022).

Machine learning is a powerful tool for detecting complex signals in various types of biological data (Eraslan et al. 2019; Vamathevan et al. 2019; Esteva et al. 2019; Wainberg et al. 2018; Xu and Jackson 2019; Ching et al. 2018; Libbrecht and Noble 2015; Greener et al. 2022). Previous studies have shown that even relatively simple ML approaches can be successfully applied to classify immune repertoires based on the immune status (repertoire-based classification), with their performance improving with dataset size (Emerson et al. 2017; Pavlović et al. 2021; Ostmeyer et al. 2019). These few promising successes showcase the potential of ML for detecting complex signals in AIRR data but many challenges remain for repertoire-based diagnostics to reach a level of maturity sufficient for clinical application (Arnaout et al. 2021). For example, although the accuracy achieved for predicting CMV status from TCR repertoires was found to be relatively high (Emerson et al. 2017), CMV is known to leave a particularly strong mark on the adaptive system with a high percentage of CD8+ T cells specific to CMV increasing with age (Kuijpers et al. 2003). To achieve clinically acceptable accuracy levels for predicting a variety of disease states, more sophisticated and sensitive ML approaches will be needed (Kanduri et al. 2021; Greiff, Yaari, and Cowell 2020; Pavlovic et al., n.d.; C. R. Weber et al. 2022; Slabodkin et al. 2023; Widrich et al. 2020; Pradier et al. 2023). Similarly to repertoire classification, it remains challenging to predict antigen binding for individual AIRs (receptor-based classification) given, for instance, the current scarcity of the antigen-labeled data, as well as challenges in quantification of generalization and negative data definition (Robert et al., n.d.; Akbar et al. 2021; Dash et al. 2017; Glanville et al. 2017; A. Weber, Born, and Rodriguez Martínez 2021; Moris et al. 2021a; Akbar, Bashour, et al. 2022; Hudson et al. 2023; Dens et al. 2023).

Progress on both repertoire and receptor-based ML classification problems is hindered by a lack of large-scale data with known ground-truth information, such as corresponding immune status and immune signal (Sandve and Greiff 2022; Pavlović et al. 2022). Such large-scale ground truth data is required for the development and benchmarking of novel AIRR-adapted ML approaches as well as the benchmarking of established approaches to test their predictive performance and capability to recover immune signals (Chen et al. 2022; Prakash, Shrikumar, and Kundaje 22--23 Nov 2022; Sandve and Greiff 2022; Romano et al. 2022; Thiyagalingam et al. 2022). Specifically, the lack of ground truth data hinders the benchmarking under various conditions such as sample size and AIRR architecture (Sandve and Greiff 2022). Therefore, it is challenging to assess using only experimental data whether an AIRR-ML method successfully learned disease-associated features or spurious factors that are associated with high predictive performance (Pavlović et al. 2022). In contrast, simulation enables the generation of AIRR datasets of virtually unconstrained size, with fine-grained control over introduced signals (ground truth) (Sandve and Greiff 2022). While simulations can bridge the gap between the lack of suitable experimental data to guide the development and benchmarking of novel AIRR-ML methods relevant for real-world applications, not all simulation approaches are created equal. A key challenge for simulations is to faithfully represent the characteristics and statistical properties of experimental AIR(R)s with high fidelity.

There exist various tools for simulating a set of individual AIRs with or without set parameters (Marcou, Mora, and Walczak 2018; C. R. Weber et al. 2020; Woodcock, Bortone, and Vincent 2020; Safonova, Lapidus, and Lill 2015; Yermanos et al. 2017; Han et al. 2022; Sutherland and Cowan, n.d.; Yang et al. 2021; Davidsen et al. 2019; Pradier et al. 2023; Isacchini et al. 2021). These tools accurately mimic the V(D)J-recombination process to varying degrees, but most do not simulate the selection or implantation of putative immune signals (C. R. Weber et al. 2020). When introducing such immune signals, it is crucial not to perturb existing and underlying statistical properties of AIRR data, such as germline gene recombination statistics (Marcou, Mora, and Walczak 2018; Slabodkin et al. 2021) and the degree of sharing of public AIR sequences between individuals (Kanduri et al., n.d.), since perturbated properties could lead to ML methods achieving high accuracy based on artificial features rather than meaningful signals.

Here, we implemented the simulation suite LIgO, which enables the generation of immune-signal-labeled synthetic AIR(R) datasets for the development and benchmarking of AIRR-based machine learning methods. Importantly, the LIgO suite allows simulating AIR(R) datasets at scales way beyond what is currently experimentally feasible. Immune signals in AIRs can be simulated based on either rejection sampling or signal implantation that preserves AIR generation probability distribution through importance sampling. LIgO can be used with immuneML, an open-source ecosystem for AIRR-ML analysis (Pavlović et al. 2021). We show the applicability of LIgO on two distinct machine learning use cases, investigating (1) how individuals carrying out-of-distribution immune signals impact receptor-based prediction performance and exploring (2) the limitations of conventional encoding schemes for AIRR binary classification when immune signals co-occur within the same AIR.

## Results

### Immune signal and immune event formalization in AIR(R) data

Adaptive immune receptors (AIRs) can be labeled with their antigen recognition capacity (one or multiple antigens) and adaptive immune receptor repertoires (AIRRs) with the individual’s disease state(s). These labels can be used to formulate a machine learning (ML) classification problem on AIRR data (Fig. 1B). Both receptor-based (AIR-based) and repertoire-based (AIRR-based) classification may be considered as a binary classification problem (the AIR binds or does not bind to the antigen), a multi-label classification problem (the AIRR is associated with multiple disease states), or a multi-class classification problem (the AIRR is associated with a specific stage of a disease, e.g., stages of cancer). As of yet, there is limited understanding of how immune signals are encoded inside AIRRs (Greiff, Yaari, and Cowell 2020). To simulate AIRR data for benchmarking receptor and repertoire-based AIRR-ML methods, there is a need to account for different types of putative immune signals that ML methods are expected to retrieve and the statistical properties of synthetic immune signals vis-à-vis AIRR-ML tasks (Fig. 1A).

**Figure 1:**
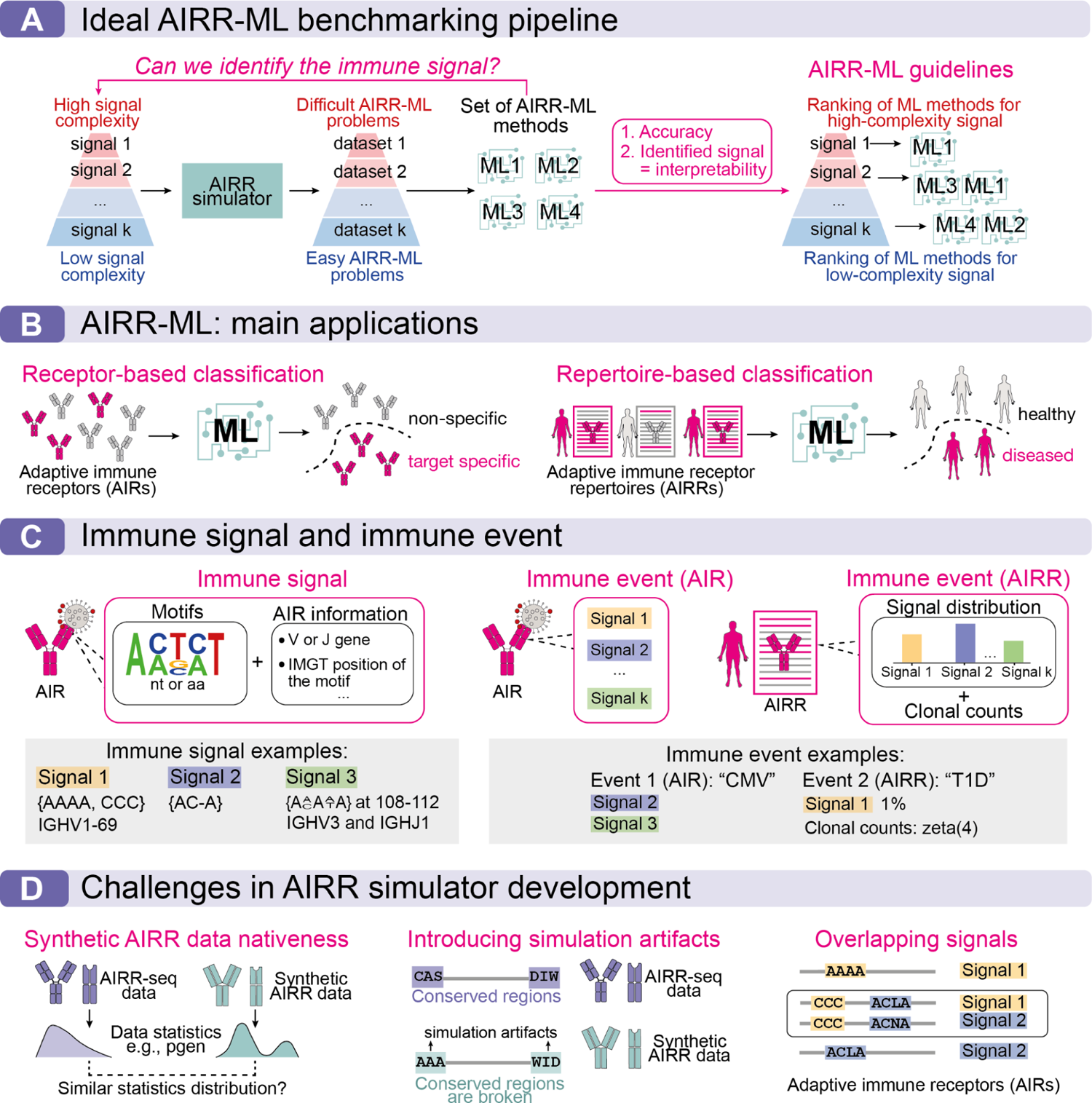
Formalization of an AIRR simulator suitable for AIRR-ML benchmarking both on the receptor and the repertoire level. (A) An ultimate goal of AIRR-ML benchmarking is to establish guidelines for achieving effective repertoire and receptor classification by taking into account the various complexities of immune signals. To achieve this goal, there is a need for ranking AIRR-ML approaches based on their prediction accuracy and ability to recover interpretable immune signals. (B) The two main classes of AIRR-ML classification problems are receptor-based and repertoire-based classification. The aim of the receptor-based classification is to predict antigen binding for individual AIRs, and the aim of the repertoire-based classification is to classify immune repertoires based on their immune status. (C) We define an immune signal as the union of a set of motifs and AIR-specific information (the latter may be absent). A motif is a distribution over nucleotides or amino acids at multiple positions. AIR-specific information can be, e.g., V or J gene, position of motif in a CDR3, signal of the other chain for the paired data. We illustrate signal definition with three different signals. Signal 1 consists of the two k-mers AAAA and CCC, and the V gene IGHV1-69, which means that only receptors with IGHV1-69 and containing either AAAA or CCC are considered signal 1 specific. Similarly to signal 1, signal 2 consists of a gapped 4-mer and signal 3 consists of a motif represented by a PWM on the 108-112 IMGT positions and restricted by IGHV3 and IGHJ1 genes. Formally, we define an immune event on the receptor and the repertoire levels. On the receptor level, an immune event is a set of immune signals. On the repertoire level, an immune event is defined as a distribution of immune signals along with clonal count distribution. Concrete examples of immune events on the receptor level may include, for instance, “cytomegalovirus infection (CMV)” which consists of signal 2 and signal 3 described above. And an AIR is considered as CMV-specific if it contains signal 2 or signal 3. We may also define an AIRR carrying a hypothetical T1D (type 1 diabetes) immune event if 1% of AIRs contain signal 1 and the clone counts of the signal-specific receptors follow a zeta distribution. (D) Developing an AIRR simulator that implements immune signal and event concepts of varying levels of complexity and is also suited for AIRR-ML benchmarking both on the repertoire and receptor level presents several challenges. (i) For real-world relevance, simulated AIRR data with implanted signals should be as similar as possible to experimental AIRR-seq data. Specifically, such similarity may be defined in terms of generation probability distribution (pgen), where pgen is the probability of a given V(D)J recombination process to generate a given AIR. (ii) A simulation process might introduce simulation artifacts into data, for example, break the conserved receptor regions. In this case, even if an AIRR-ML approach perfectly detects the implanted signal, it will differ from biologically relevant signals. (iii) One receptor might contain multiple signals corresponding to different immune events and thus immune signals and immune events can be overlapping both on the receptor and repertoire levels which complicates their simulation.

To support a broad variety of potentially interesting AIRR-ML problem formulations, we opted for a very flexible, yet structured, specification of immune signals. We decided to separate between biological immune events — e.g., disease, allergy, vaccination, or other situations that elicit an adaptive immune response — and associated encoded signals in the receptors. The AIRs produced during the immune response and associated with a (biological) immune event are considered immune-event specific. Each immune-event specific receptor contains one or more immune signals, which, for instance, reflects its binding rules to the immune event antigen(s) (Fig. 1C). To prevent excessive complexity, we limit the number of immune signals per receptor, ensuring that each receptor contains no more than two immune signals.

We define that on the receptor level, an *immune signal* consists of a set of *motifs*, where each motif is a distribution over amino acids or nucleotides of any length up to an entire AIR sequence (Emerson et al. 2017). Furthermore, additional AIR information is defined, such as V and J genes, and the distribution of the motif set locations within the CDR3 (Fig. 1C), in line with previous studies (Emerson et al. 2017; Kanduri et al. 2022; Ostmeyer et al. 2019; Akbar et al. 2021). In addition, AIR information may include derived features such as CDR3 length (Shemesh et al. 2021), physicochemical properties (Thomas et al. 2014), or binding energy (Robert et al. 2022). In the case of paired chain data, the immune signal would include this information from both chains (Akbar et al. 2021; Jaffe et al. 2022; Glanville et al. 2017; Dash et al. 2017).

On the receptor level, we define an *immune event* as a receptor label associated with a set of immune signals (Fig. 1C). For example, an HIV immune event may contain several immune signals corresponding to different HIV epitopes. If an AIR is polyreactive, it is linked to multiple immune events. If an AIR is not specific to any annotated antigen, then it has no label.

On the repertoire level, we define an immune event as a distribution of immune signals in conjunction with a distribution of clone counts (Fig. 1C), which can represent, for example, clonal expansion (Greiff et al. 2015). Our definition of immune signals and events on the receptor and repertoire levels is designed to encompass a wide range of data formats and biological signals in a comprehensive manner that have been previously reported on experimental data (Greiff, Yaari, and Cowell 2020). From here on, we will not separate the terms immune event and immune signal on the repertoire and receptor level.

### Simulation approaches for generating biologically relevant AIR(R) data

To fit the AIRR-ML problem formulation, one needs to simulate signal-specific (positive) AIRs and non-specific (negative) AIRs and control their frequencies in an AIRR dataset to encode an immune event. In this study, we consider two approaches to simulate biologically relevant AIRs carrying immune signals — rejection sampling and signal implantation (Fig. 2C). With rejection sampling, a pool of random background AIRs is generated and only the AIRs containing a predefined signal(s) are considered signal-specific. The main advantage of the rejection sampling approach is that it does not modify the original AIRs, meaning that receptor-based statistics like generation probability for each individual AIR are not artificially perturbed (Fig. 1D). However, rejection sampling is not computationally feasible for immune signals with low population probability, since estimated time of finding an immune-signal-specific receptor in background receptors is inversely proportional to its population probability and large number of AIRs needs to be generated for each receptor that is kept after signal filtering. An alternative way to generate signal-specific receptors is signal implantation, which replaces a substring of a CDR3 region of a given background AIR with one of the immune signal (gapped) k-mer, or motif in general. Unlike rejection sampling, computational efficiency of signal implantation does not depend on the population probability of the immune signal. However, signal implantation may introduce artifacts to simulated AIRs because it modifies the original sequence and may create unrealistic sequences (Fig. 1D). Also, signal implantation does not work for every immune signal, since some signals cannot be easily implanted when an AIR is already constructed - e.g., if the signal to be implanted is a specific V gene, then substituting the original V gene with the new one might potentially conflict with the CDR3 sequence of the original AIR. Both rejection sampling and signal implantation may be coupled with an optional importance sampling step. Importance sampling minimizes potential perturbation of simulated AIR pgen distribution, which, if left unchecked, may cause artificial increase of ML performance. Depending on the defined signals, LIgO will allow testing whether rejection sampling or implantation methods can be performed. Altogether, both the rejection sampling and the implantation approach allow generation of datasets with a controlled number of signal-specific receptors in the synthetic data.

**Figure 2:**
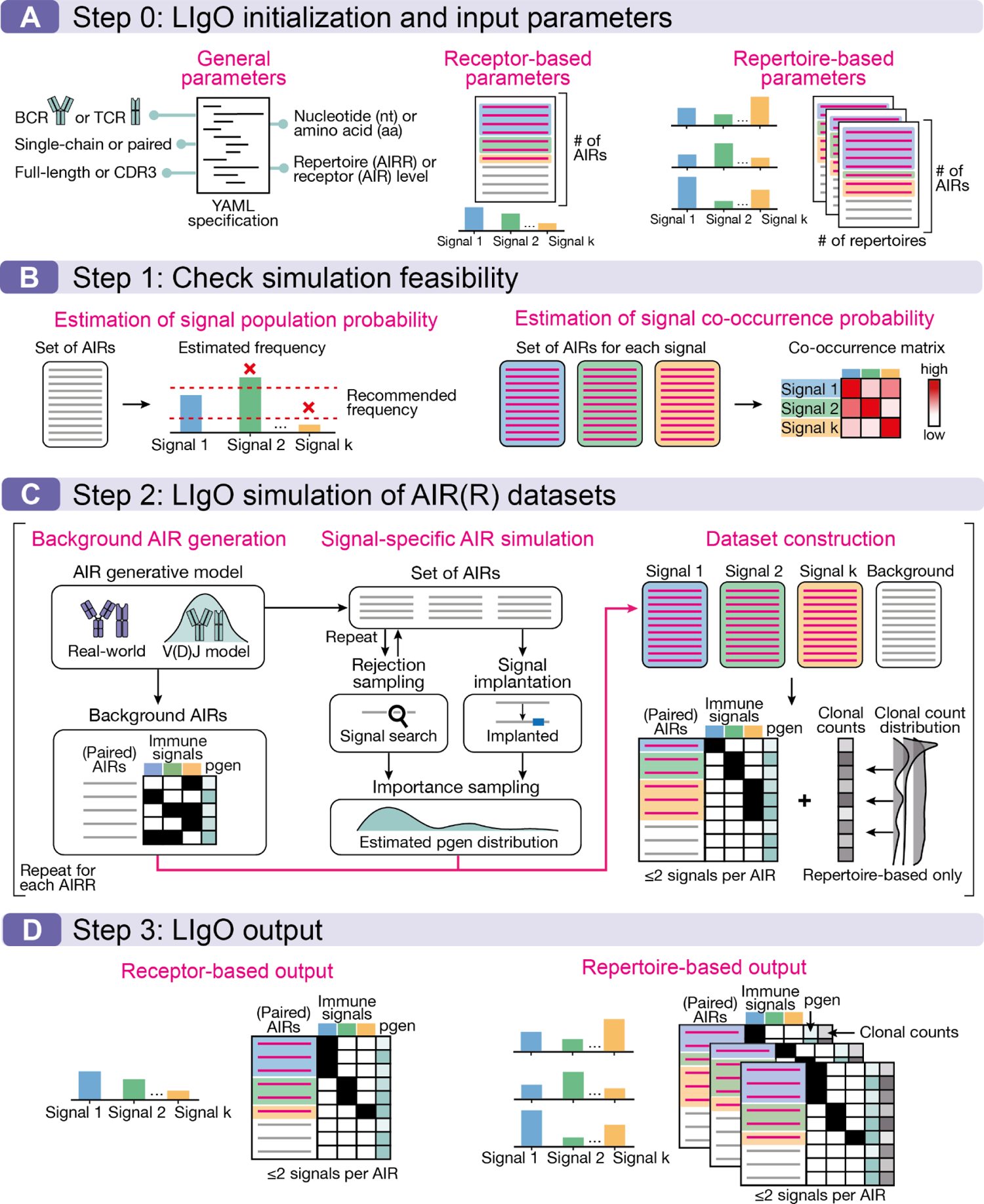
LIgO workflow for configuring AIRR-ML benchmarking datasets. (A) For initialization, a user may define general simulation parameters, such as the type of AIR (BCR or TCR), receptor- or repertoire-based simulation, paired- or single-chain receptors, type of sequence alphabet — nucleotide (nt) or amino acid (aa), output format — full-length (Heather et al. 2022) or CDR3 only. For the receptor-based simulation, a number of generated AIRs and a set of immune signals with their frequencies in the final dataset are defined. For the repertoire-based simulation, a number of repertoires, number of receptors in a repertoire, and a mixture of immune signals corresponding to each repertoire with their frequencies are defined. In total, the sum of all immune signal frequencies including co-occurring signal frequencies for each AIRR should be less or equal to 1. (B) The optional second step helps users to check simulation feasibility based on the parameters defined during initialization. The first helper function estimates the frequency of each immune signal in a set of AIRs. It is not recommended to use rejection sampling for immune events whose estimated frequency falls below the recommended lower bound, by default 0.001%. In addition, the user can redefine the immune signal with low frequency in order to increase its frequency, e.g., simplify the immune signal definition by removing certain V gene(s) or by simplifying the motif in the definition. It is also not recommended to use immune signals with estimated frequency above the upper bound of 10% because they may impact the biological nativeness of immune signals given the rarity of such high-frequency signals. The second helper function estimates the co-occurrence of defined immune signals. In order to limit computational complexity, LIgO restricts the number of immune signals per one AIR to at most two and all AIRs containing three or more immune signals will be removed during the simulation process. If the user wants to limit the number of immune signals per AIR by one, then immune signals frequently co-occurring in the background may decrease rejection sampling efficiency. It is recommended for the user to redefine immune signals in this case. (C) In the main simulation step, LIgO requires a generative model to obtain background AIRs, which can either be simulated from a V(D)J-recombination process (e.g., IGoR (Marcou, Mora, and Walczak 2018; Sethna et al. 2019) or immuneSIM (C. R. Weber et al. 2020), where a custom V(D)J model can be inferred from real-world data) or taken from experimental data (e.g., iReceptor (Corrie et al. 2018) or OAS (Kovaltsuk et al. 2018)). Subsets of immune-signal specific receptors can be simulated using rejection sampling or implantation from a set of AIRs which may or may not have the same V(D)J generation parameters as the background AIRs. Additionally, if the simulation is defined by a V(D)J model, the immune-signal specific AIRs can be subsampled according to their generation probabilities during the importance sampling step. Then, background and immune-signal-specific AIRs are combined in one dataset with respect to all immune signals, and AIRs containing more than two immune signals are removed. For repertoire-based simulation, clonal counts are sampled from a zeta distribution (Greiff et al. 2015) with user-defined parameters for each immune event or constructed based on the normalized receptor generation probabilities. If the simulation is performed on the repertoire level, then the main simulation step should be repeated *n* times, where *n* is the number of desired AIRRs to be simulated. (D) LIgO reports all parameters used during the simulation steps. For the receptor level simulation, a set of (paired) AIRs (V and J genes, CDR3 region and optionally the full-length receptor) is reported together with their specificity to each immune signal and immune event, generation probability (optional), and immune signal position in each AIR. For the repertoire level simulation, receptors of each repertoire are annotated with the immune signal specificity matrix, immune signal position in each AIR, generation probability of each AIR (optional), and clonal counts. Repertoire-level metadata with immune signal and immune event annotations is also available in a metadata file accompanying the dataset. LIgO outputs datasets in AIRR-compliant format both for receptor and repertoire-level simulations.

### LIgO simulation workflow

Here we present the LIgO workflow that enables simulation of immune receptor (AIR) and repertoire (AIRR) data for the development and benchmarking of AIRR-based machine learning. As the first step (Fig. 2B), LIgO estimates the feasibility of a simulation described with a given set of user-defined simulation parameters (Fig. 2A) by assessing two potential bottlenecks (Fig. 2B). First, LIgO determines if the user-defined signals can be efficiently generated using rejection sampling. If this is not the case, LIgO suggests using signal implantation optionally coupled with importance sampling or modifying the signal(s) definition, because attempting to simulate rare immune events using rejection sampling can lead to the first computational bottleneck. The second bottleneck may arise if many AIRs violate our complexity restriction, which specifies that each AIR must contain no more than two immune signals. In such cases, these AIRs will be eliminated from the simulation process which may also increase simulation time. LIgO assesses if any two immune signals frequently co-occur in the same AIR and suggests the user to redefine co-occurring immune signals. Both bottlenecks can notably increase the simulation time and may render it quasi-infinite. Although the feasibility assessment step is optional, we recommend users to check the simulation feasibility before running the main simulation step.

LIgO initiates the main simulation step by obtaining background receptors to be used as a basis for simulation (Fig. 2C). It does so by either utilizing a generative model, which offers the flexibility of, for instance, using a custom IGoR V(D)J model (Marcou, Mora, and Walczak 2018; Sethna et al. 2019; Slabodkin et al. 2021) or an experimental AIRR-seq dataset. While the V(D)J model can simulate an unlimited number of AIRs, it may contain simulation artifacts. Experimental AIRR-seq data might be more realistic, but they are of a limited size and may contain unknown artifacts (Pavlović et al. 2022), such as batch effects and AIRR imprinting of previous diseases, (Sandve and Greiff 2022), thus potentially rendering them insufficient for complex LIgO simulations.

Once the background receptor dataset has been obtained, LIgO achieves the desired amount of immune-signal-specific AIRs using rejection sampling and signal implantation (Fig. 2C). To enhance the rejection sampling efficiency for immune signals involving specific V or J genes when using a generative model to obtain the background receptors, LIgO constructs a skewed V(D)J model. This specialized model generates AIRs containing only the required V or J genes, automatically meeting the gene criteria for the given immune signal. If the simulation is defined by a V(D)J model, the user can evaluate the generation probability (pgen) of each AIR during the simulation process. Although pgen evaluation is time-consuming, the generation probability distribution can be used to further filter signal-specific AIRs during the importance sampling step (Fig. 3E). Importance sampling controls the overall pgen distribution and provides higher chances for signal-specific AIRs with more likely pgens of being included in the final dataset. If the user wishes to add background receptors in the resulting AIR(R) dataset(s), LIgO provides background receptors generated by the V(D)J model or experimental data. Additionally, they can be further filtered so they are devoid of immune signals. If the filtration is not performed, immune signals in the background receptors may impact the overall immune signal frequencies in the final simulated dataset(s).

**Figure 3:**
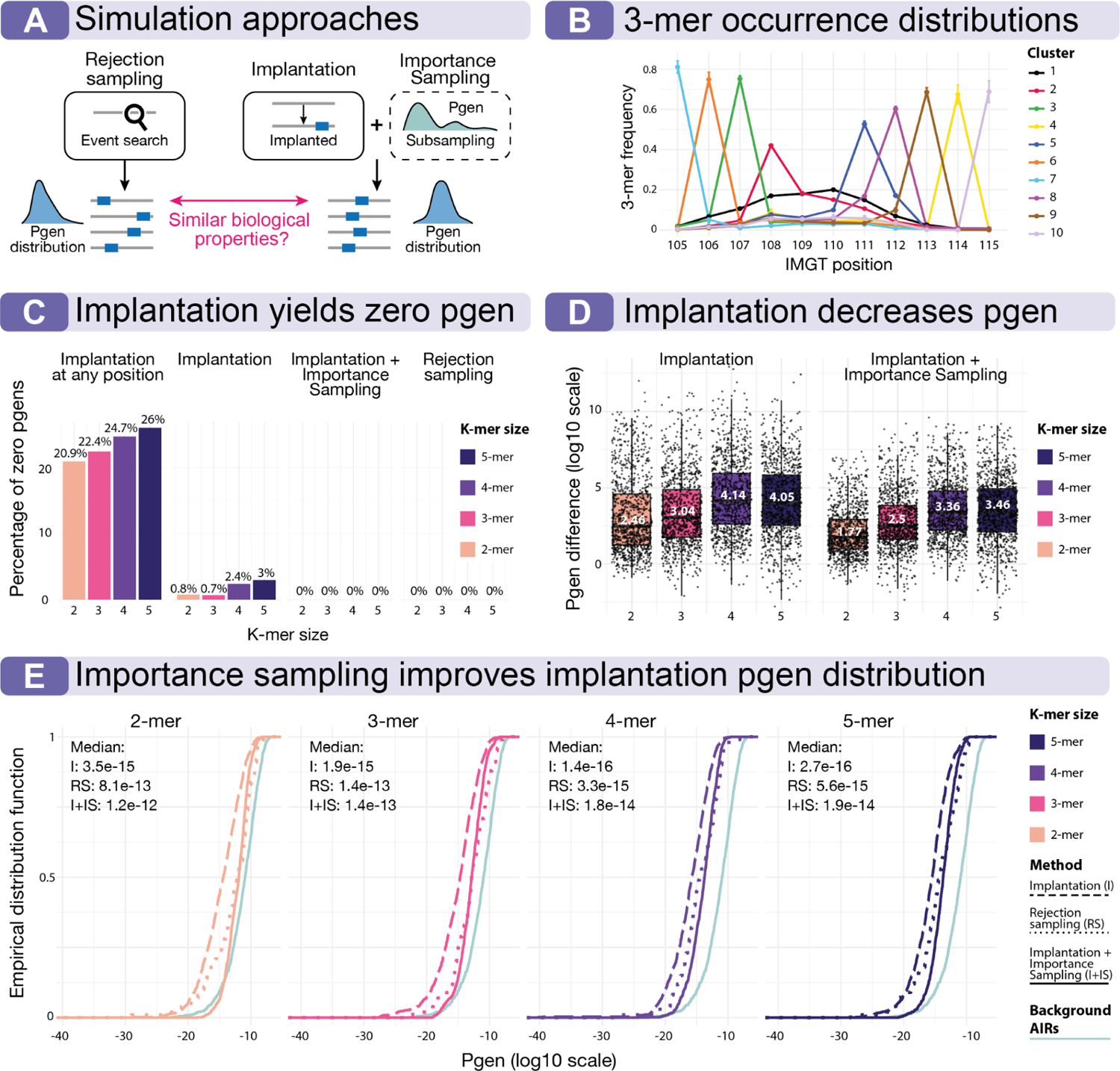
Comparison of biological properties of different simulation strategies. (A) We compared three simulation strategies — (i) rejection sampling, which keeps an AIR sequence but may be computationally inefficient, (ii) signal implantation, which changes an AIR sequence and thus may change its pgen, but is computationally efficient, and (iii) signal implantation combined with importance sampling which preserves the pgen distribution of simulated AIRs and is computationally efficient. (B) We illustrate that different immune signals may have distinct occurrence distribution across CDR3 positions, where positions are IMGT-standardized. Here we show position distribution patterns of 3-mers occurring in amino acid TRB sequences generated using the default IGoR model (Marcou, Mora, and Walczak 2018). Position frequencies were clustered into 10 clusters using hierarchical clustering, and one line on the plot represents the mean frequency for each cluster. Some 3-mers have one dominating position with occurrence probability above 0.4 while others 3-mers occurrence distribution is more uniform (cluster 1 shown in black). (C) Implantation at any CDR3 position and implantation in non-conserved CDR3 regions (shown as implantation) yields AIRs with pgens=0, which therefore can not be generated by the VDJ recombination model. However, when implantation is coupled with importance sampling, it prevents the generation of impossible AIRs, aligning with rejection sampling results. (D) Both implantation and implantation coupled with importance sampling decrease on average the pgen of the simulated AIR compared to the pgen of the original AIR before implanting. Each boxplot value corresponds to one AIR and represents log10 difference of its pgen before and after signal implantation. Implantation followed by importance sampling has lower median log10 pgen difference values compared to implantation alone, the median value for each k-mer size is shown in white. (E) The pgen distribution of signal-specific AIRs generated using implantation (shown with dashed line) is shifted compared to the pgen distribution of signal-specific AIRs generated through rejection sampling (shown in dotted line). However, this shift may be corrected using the importance sampling strategy (shown in solid line). The turquoise solid line illustrates the pgen distribution of the background AIRs, which were used as the source of AIRs for all simulation strategies.

Upon completion of the simulation, LIgO constructs AIR(R) dataset(s) with the desired frequencies of immune-signal-specific receptors for each immune event. In the scenario of repertoire-based simulation, vectors of simulated clonal frequencies can be optionally added to each AIRR. By adjusting the parameters of the clonal frequency distribution for each immune signal, users have the capability to simulate processes resembling affinity maturation and the expansion of signal-specific clones. LIgO outputs AIRR-compliant data, ensuring that each receptor is productive and contains no more than two immune signals (Fig. 2E). Each AIR can be outputted in a full-length (Heather et al. 2022) or CDR3 only format. Additionally, to facilitate further ML processing, the data is annotated with a binary matrix showing for every immune signal its presence in every receptor and, if so, the positions in the receptor sequence.

### Comparison of simulation strategies implemented in the LIgO simulation workflow

As discussed earlier, LIgO simulation approaches have their advantages and disadvantages. Rejection sampling maintains the AIR sequence unchanged but lacks computational efficiency compared to signal implantation. We investigated how one can use signal implantation to attain results similar to those obtained through rejection sampling (Fig. 3A). First, if an immune signal is defined as a k-mer or more generally as a motif, then the user should take into account the natural signal occurrence distribution across CDR3 positions and implant the motif according to the natural distribution. For a 3-mer amino acid signal, only approximately half of the motifs were evenly distributed across the CDR3, while the other half tended to occur on one CDR3 position (Fig. 3B, Supplementary Fig. 1). Second, if an immune signal is implanted into any CDR3 position, then around 20% of all signal-specific AIRs will have zero pgen after implantation, which means that these zero-pgen AIRs cannot be generated through the VDJ recombination model. If we restrict the implantation position and do not implant in the first and the last two CDR3 positions corresponding to conserved CDR3 regions, then the percentage of zero-pgen AIRs drops to around 2% (Fig. 3C). In general, signal implantation decreases pgen of the original AIR by two to four orders of magnitude on average (Fig. 3D). However, when signal implantation is coupled with importance sampling, LIgO will no longer produce signal-specific AIRs with zero pgen values, and the overall pgen difference after implantation is less pronounced than with pure signal implantation (Fig. 3C, Fig. 3D). Finally, we demonstrated that by employing signal implantation coupled with importance sampling to simulate signal-specific AIRs, we can achieve an overall similar pgen distribution as those generated through rejection sampling, while not increasing the computational time (Fig. 3E).

### Use case 1: Out-of-distribution receptor-level simulation using LIgO

Train and test data composition may impact AIRR-ML models’ quality and accuracy (Gygi, Kleinstein, and Guan 2023). For example, (i) if the train and test set overlap or contain highly similar sequences, then the accuracy of the trained ML model may be overly optimistic (a case of data leakage). (ii) If the training data is not comprehensive and representative, it may lead to generalization problems and underperformance on unseen data (Montemurro, Jessen, and Nielsen 2022; Deng et al. 2023; Dens et al. 2023; Moris et al. 2021a; Robert et al. 2022; Meysman et al. 2023; Walsh, Pollastri, and Tosatto 2016; Petti and Eddy 2022; A. Weber, Born, and Rodriguez Martínez 2021). Since different individuals have largely non-overlapping AIRRs (Warren et al. 2011; Briney et al. 2019) and, thus, potentially different immune signals to the same targets, we investigated how immune signal variability may impact ML accuracy and model generalization to other unseen (potentially out-of-distribution) individuals.

For this use case, we considered a scenario where an immune signal is defined as one motif, yet each individual carries a slightly different modification of this motif. Specifically, we defined the immune signal as a motif AA-A with four variations of the immune signal — AAAA, AANA, AACA, AAGA, reflecting AIRs from four different individuals (Fig. 4B). We simulated 4⋅10^4^ AIRs and replicated the experiment ten times, which resulted in 4⋅10^5^ of simulated AIRs in total. Simulation details can be found in the Methods section. We illustrated this use case with a binary receptor-based classification task (Fig. 4A), categorizing AIRs into signal-specific (i.e., an AIR contains one out of four signal 4-mers) or non-specific (i.e., an AIR does not contain any signal 4-mers). We trained and evaluated a logistic regression (LR) model using two train-test strategies — (i) random, which randomly splits all receptors from four individuals into train and test set, and (ii) leave-one-individual out, which places all AIRs from one individual into a test set and AIRs from all the other individuals into a training test (Fig. 4C).

**Figure 4:**
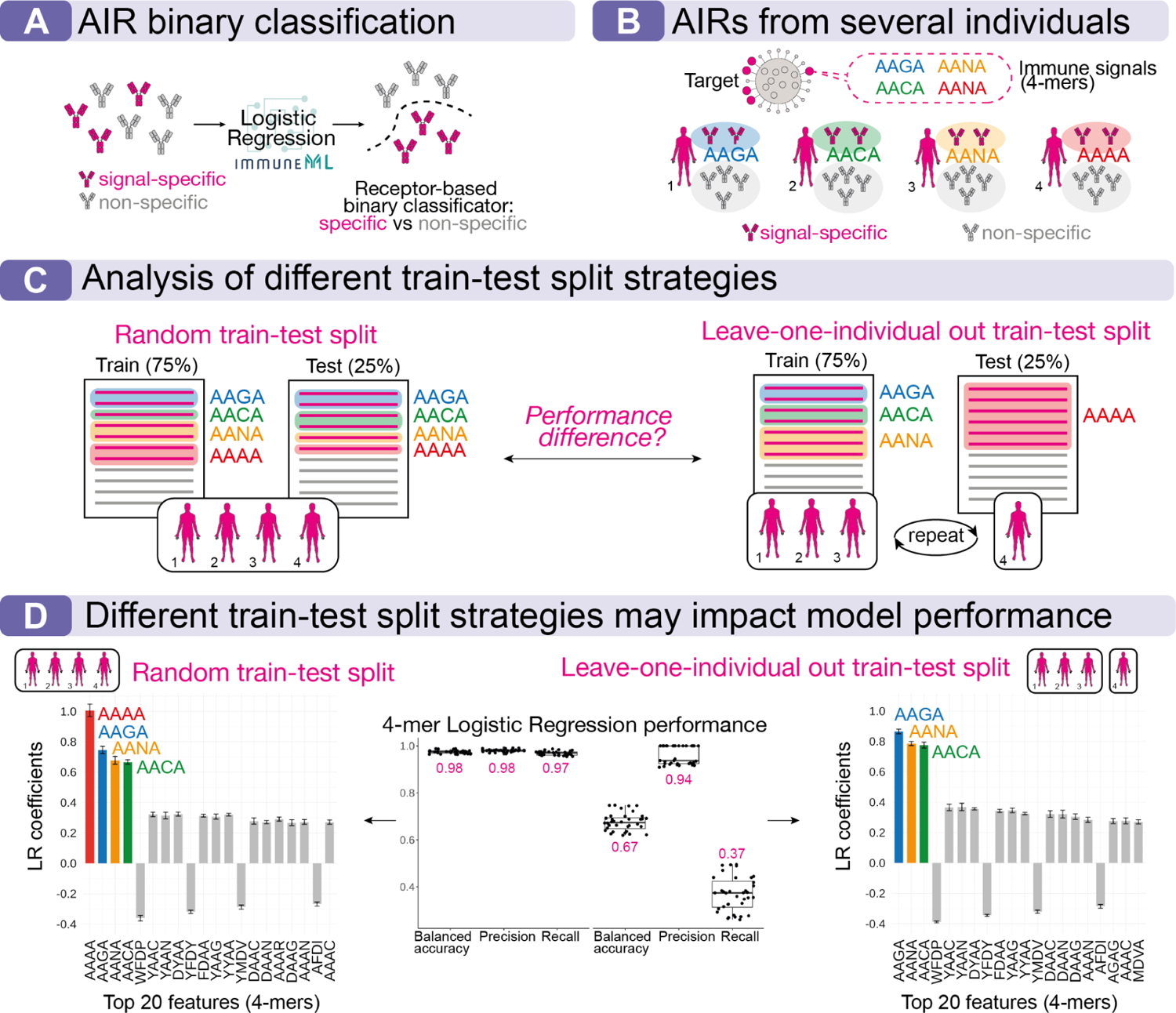
Use case 1: Out-of-distribution receptor-level simulation using LigO. (A) We illustrate an example of an out-of-distribution problem using logistic regression as a ML method for receptor-based binary classification, i.e., classification of signal-specific vs non-specific AIRs. See Methods section for more details. (B) We defined the immune signal by four 4-mers differing in the third position — AAGA, AANA, AACA, AANA. We simulated four sets of 10000 AIRs using signal implantation, where 5000 AIRs were signal-specific, and 5000 AIRs were non-specific, i.e., not containing any signal 4-mers. We assumed that one set of AIRs corresponds to one individual, and each of the four sets of AIRs contained only one out of four signal 4-mers. We replicated the simulation ten times. (C) We applied two train-test split strategies to train the logistic regression model for receptor-level classification — (i) random split, where the signal-specific AIRs containing the four signal 4-mers are present both in the train and in the test set, and (ii) leave-one-individual-out split for every individual, where AIRs with only three 4-mers out of four were present in the train data and the AIRs with the fourth signal 4-mer was present in the test data only. (D) Performance comparison and interpretability analysis. We trained a 4-mer logistic regression with two train-test split strategies on all ten replicates resulting in ten ML models. The 4-mer logistic regressions trained on the random train-test split achieved higher accuracy and its largest coefficients corresponded to the four signal 4-mers (AAAA, AAGA, AANA, AACA). The 4-mer logistic regression trained on the leave-one individual out train-test split achieved lower performance with a median balanced accuracy of 0.57. The LR achieving the best performance was trained on the split, where all AIRs containing the 4-mer AAAA were placed in the test set. The largest coefficients of this LR contain three out of four signal-specific 4-mers (AAGA, AANA, AACA). The boxplots display model performances for all four splits within 10 replicates and the median performance value is shown in red. The bar plots display the top twenty logistic regression coefficients from the best model with the highest absolute value, and the error bars indicate the standard deviation for ten replicates.

The LR trained on the random train-test strategy achieved higher balanced accuracy than the LR trained on the leave-one-individual-out train-test approach (Fig. 4D). Based on interpretability analysis, it can be seen that the random train-test model obtained and learned information about all 4-mers while the leave-one-individual out model did not learn the signal that was not present in the training set. This use case illustrates the impact of train and test data on AIRR-ML quality. It demonstrates that while perfect prediction accuracy may be observed with a random split approach, prediction accuracy may come across as more limited based on a leave-one-individual-out split strategy. Leave-one-individual-out split strategy is considered more real-world relevant since it is likely that a limited number of individuals does not suffice to cover the entire antigen-specific AIR sequence diversity.

### Use case 2: Limitations of conventional encoding schemes for repertoire-level binary classification when immune signals co-occur within the same AIR

Several previous studies have suggested that the information needed for repertoire-based classification is sufficiently represented through conventional k-mer encoding of AIRs (Akbar et al. 2021; Ostmeyer et al. 2019; Katayama and Kobayashi 2022; Cinelli et al. 2017; Sun et al. 2017; Thomas et al. 2014). However, immune signals may comprise multiple co-occurring motifs within the same receptor (Dash et al. 2017; Glanville et al. 2017; Akbar et al. 2021). In this case, conventional k-mer encoding that treats each k-mer in isolation may not be sufficient to represent the distinction between immune states. LIgO can simulate such complex immune signals that may pose challenges to ML methods that do not consider the co-occurrence of motifs within the same receptor. To demonstrate this, we simulated AIRR datasets for a repertoire classification problem, where the distinction between contrasting immune states cannot be observed using conventional k-mer encoding (Fig. 5A).

**Figure 5:**
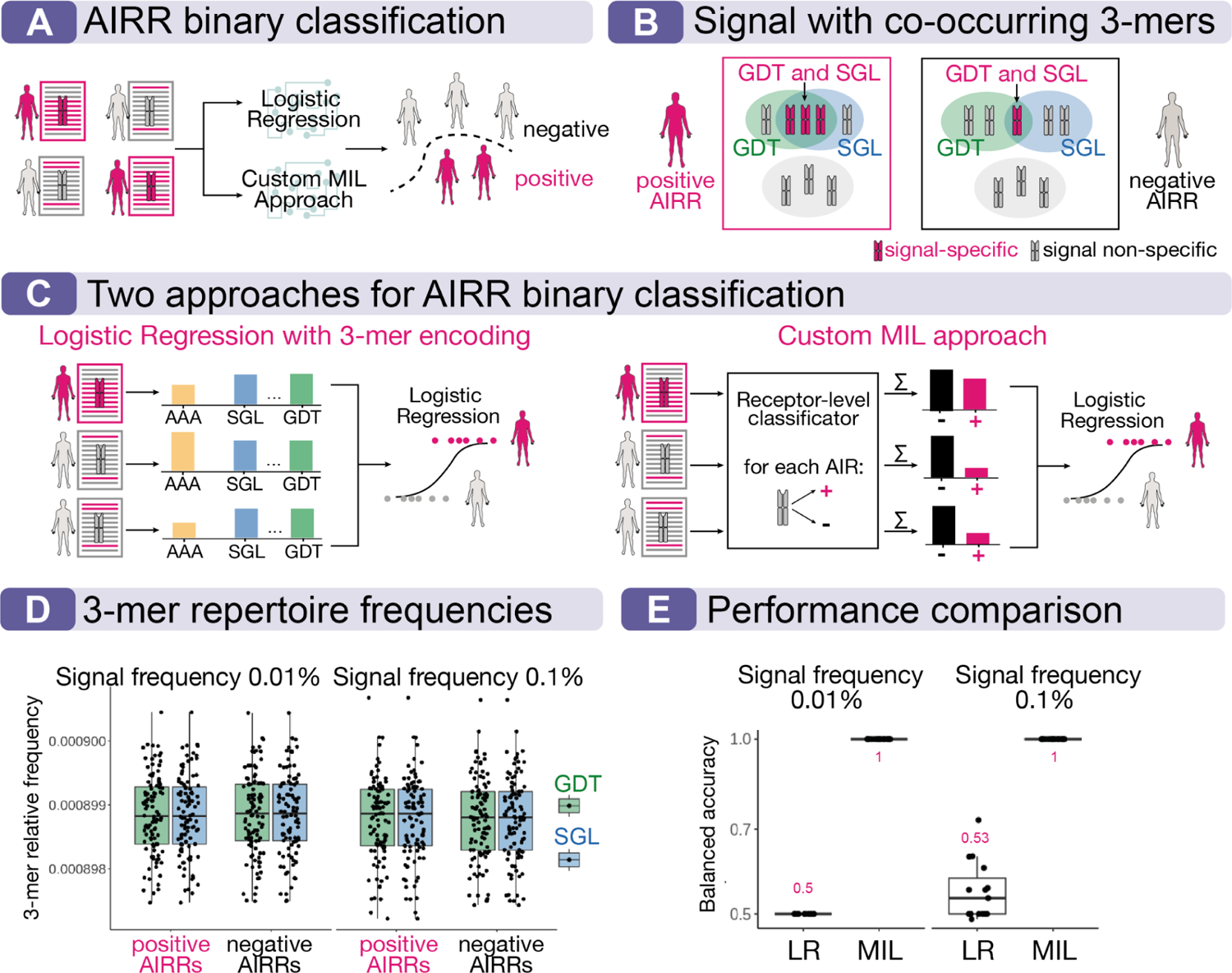
Use case 2: Limitations of the conventional k-mer encoding scheme for repertoire-level binary classification when immune signals co-occur within the same AIR. (A) We demonstrate the limitations of conventional k-mer encoding for a binary AIRR classification problem by utilizing LR and a custom multiple instance learning (MIL) approach. The task involves classifying repertoires as positive or negative based on the presence of an immune signal. Detailed methods can be found in the Methods section. (B) An example of a complex immune signal that may pose challenges to ML methods that neglect the co-occurrence of immune signals within the same AIR. The signal is composed of co-occurring GDT and SGL 3-mers within the same AIR and the frequency of such AIRs is higher in the positive repertoires. Moreover, the frequency of each 3-mer GDT and SGL remains identical in both the positive and negative repertoires, rendering it indistinguishable using the conventional k-mer encoding approach. We simulated two scenarios where positive repertoires consist of 0.01% and 0.1% of signal-specific receptors containing both GDT and SGL. For each scenario, we simulated 200 TRB repertoires (100 AIRRs in each class and 10^5^ AIRs per repertoire) and replicated the simulation three times (1200 AIRRs and 1.2⋅10^8^ TRBs in total). (C) We tested two MIL approaches for binary repertoire classification — (i) LR with conventional 3-mer encoding and a custom MIL approach. Briefly, the conventional LR approach (i) encodes every repertoire to a vector of normalized 3-mer frequencies across all AIRs within the repertoire. After that, vectors of 3-mers are classified using LR. The second custom MIL approach (ii) takes into account the 3-mer co-occurrence on the receptor level. First, every receptor is classified into signal-specific and non-specific based on the Fisher’s exact test. On the second step, a LR classifies repertoires based on the number of signal-positive and negative receptors within the repertoire. (D) The relative frequency distribution (on the y-axis) of two 3-mers, GDT and SGL, that constitute the immune signal is shown for positive and negative repertoires and two different immune signal frequencies (0.01% and 0.1%). Note that the y-axis here represents normalized frequencies relative to the frequencies of all k-mers and is different from the immune signal frequency percentages. The visualization shows that the frequencies of the immune signal 3-mers is similar for positive and negative AIRRs when their joint co-occurrence is not considered. (E) Performance estimates (on the y-axis) of two different ML methods in a binary repertoire classification shown with balanced accuracy. As expected, the conventional LR with a 3-mer encoding performs poorly, achieving median balanced accuracy of 0.5 and 0.53 in two signal frequency scenarios. In contrast, the custom MIL, which considers the 3-mer co-occurrence, achieves a median balanced accuracy of 1 in both signal frequency scenarios.

Specifically, using LIgO, we simulated many TRB repertoire datasets, where each dataset contained 200 repertoires (100 each of positive and negative-labelled) with 10^5^ productive CDR3 amino acid sequences per repertoire. Simulation details are described in the Methods section. We assumed that the immune signal that differentiates positive and negative-labelled repertoires is defined as the joint co-occurrence of two amino acid 3-mers: GDT and SGL (Fig. 5B). The experiments were performed in two different settings with varying frequencies at which the immune signal occurs: 0.01% and 0.1%. The simulation with LIgO was carried out in such a way that the relative abundances of the chosen 3-mers, when considered in isolation, is similar for both positive and negative-labelled repertoires (Fig. 5D). The distinction between the labels will only be apparent when the receptor-level co-occurrence of GDT and SGL is considered.

We assessed the performance of the logistic regression (LR) with the conventional k-mer encoding on the LIgO-simulated data with co-occurring immune signals. In our previous study (Figure 3.a in (Kanduri et al. 2022)), we showed that the LR approach exhibited close to perfect performance even at signal frequency of 0.01% if a signal is defined as a set of motifs and each AIR contains no more than one signal motif. However, the same method exhibited poor performance even at a very high signal frequency of 0.1% in the current simulations where the signal is defined as two co-occurring 3-mers (Fig. 5E). As a proof of concept, we implemented a custom multiple instance learning approach (MIL) which is based on the underlying assumption that the immune signal is encoded in 3-mer pairs. The detailed description of the custom MIL approach can be found in the Methods section (Fig. 5C). In contrast to the LR with the conventional k-mer encoding, the custom MIL approach was able to perfectly classify the repertoires even at a signal frequency of 0.01% (Fig. 5E) with median balanced accuracy of 1. Given that the simulation design did not include any label or attribute noise, and that the immune signal frequency is quite distinct between positive and negative labelled repertoires (∼ 10 AIRs vs ∼ 1 AIR harbouring signal), it was expected that the ML model exhibited near-perfect performance. To summarize, benchmarking the standard k-mer ML approach for repertoire classification with LIgO simulated data shows that they are unable to detect co-occurring immune signals. This demonstrates how LIgO can be used to assess the capacity and limits of specific approaches.

## Discussion

Simulations, as previously emphasized by Sandve and Greiff (Sandve and Greiff 2022), enable the benchmarking of computational methods and are valuable in bioinformatics research, even in the presence of sufficient experimental data with known ground truth. Here, we developed the LIgO software suite for flexible simulation of AIRR data suitable for AIRR-ML benchmarking. While there exist other software suites for AIRR data simulation (Marcou, Mora, and Walczak 2018; C. R. Weber et al. 2020; Safonova, Lapidus, and Lill 2015; Sethna et al. 2019), these tools do not offer the possibility for configuring entire AIRR-ML benchmarking datasets for both repertoire and receptor-based classification tasks with complex and native-like immune signals (Supplementary Table 1). LIgO can be used together with the immuneML framework (Pavlović et al. 2021), enabling reproducible integration of simulation and subsequent AIRR-ML experiments (Supplementary Fig. 3).

To illustrate the usefulness of simulations for AIRR-ML research, we demonstrated LIgO on two use cases — (i) how train and test data split may drop the observed predictive performance in the out-of-distribution case, and (ii) how sequence encoding schemes can capture varying degrees of immune signal complexity. Using the ground truth information about immune signals present in LIgO-simulated AIRR data, we compared the immune information captured by the ML approaches to the original immune signals present in the data. The first use case demonstrated that the difference in underlying signal distribution between train and test sets may impact the ML method’s observed performance. The second use case shows how LIgO simulations can be set up to reveal how a custom MIL method built based on a data assumption, like assuming that antigen specificity is determined by a pair of signals, indeed outperforms a standard approach if applied to data following this assumption. These use cases illustrate how LIgO can be used to investigate connections between the assumptions used for building an AIRR-ML method and the assumptions made about the AIRR data. These use cases are not yet feasible to reproduce on experimental data since ground truth information about immune signals present in every AIR is usually not known (Akbar, Bashour, et al. 2022; Robert et al. 2022). The two demonstrated use cases are only example usages of the LIgO package. In addition, LIgO can also be utilized for benchmarking other AIRR methods, such as evaluating the performance of antigen-specific AIR clustering approaches (Valkiers et al. 2021; Rognes et al. 2022; Chang et al. 2023).

A main difficulty with AIRR simulations suitable for AIRR-ML benchmarking is the poorly described nature of immune signals. Although it is widely agreed upon that immune signals exist on the frequency level (clonal expansion) (Greiff et al. 2015), sequence level (binding and structure-related sequence motifs) (Ostmeyer et al. 2019; Akbar et al. 2021), and structure level (Akbar et al. 2021), their antigen- or disease-specific shape remains largely uncharacterized. Therefore, simulation frameworks that aim to enable real-world relevant ML benchmarking need to encompass a large array of potential immune signal definitions and configurations to cover a broad area of the potential biological immune signal space. To this end, LIgO immune signals may be defined through gapped k-mers or motifs in general (Greiff, Weber, et al. 2017; Ostmeyer et al. 2019), repertoire generation model (Slabodkin et al. 2021), clonal expansion (Greiff et al. 2015), or a combination thereof. Additionally, LIgO offers the flexibility for immune signals to take any shape when defined through a custom function that indicates the signal’s presence in an AIR. Such functionality was lacking in previous simulation tools (Supplementary Table 1). Although LIgO offers limitless possibilities for defining immune signals, it remains the user’s responsibility to define an immune signal that comprehensively captures biological complexity and reflects underlying assumptions of an AIRR-ML model to be validated. Particular care is taken to ensure that immune signals do not break underlying AIR biology, such as AIR germline-originating subsequences and the generation probability distribution. We demonstrated that signal implantation coupled with importance sampling may achieve a similar pgen distribution to rejection sampling while keeping computational efficiency. The flexibility of the presented approach allows users to define immune signals according to the state of understanding in the field at any point in time. Furthermore, helper functions guide the user as to which signal definitions are possible within a reasonable simulation time window.

In the future, large-scale antigen-specific AIRR data both on the receptor and repertoire level will be necessary to understand immune signals better. More specifically, on the repertoire-level, the dynamics of repertoire structure as a function of antigen exposure require further investigation (Abu-Shmais et al. 2023). On the receptor level, we need to better understand the long-range dependencies that govern paratope-epitope interaction, using structural (Akbar et al. 2021) and deep-mutational scanning screens (Mason et al. 2021). Although LIgO does neither leverage HLA-background information for TCR repertoire simulation (DeWitt et al. 2018) nor the influence of IG/TRB germline gene background on AIRR diversity (Pennell et al. 2023; Rodriguez et al. 2023; Omer et al. 2022; Peres et al. 2023; Slabodkin et al. 2021), such influence may be modeled similarly to immune states with respect to the underlying causal model as recently shown by Pavlović and colleagues (Pavlović et al. 2022).

While LIgO encompasses many necessary features for comprehensive and exhaustive AIRR-ML simulation, possible improvements are: (i) integrate structure-based features enabling implicit and nature-like inclusion of long-range dependency signals (Robert et al. 2022). (ii) Combine LigO with sequencing read simulators (Huang et al. 2012; Gourlé et al. 2019) to incorporate PCR or sequencing errors. This type of error simulation is essential for studying artificially induced diversity versus biological clonal expansion and batch effects (Smirnova et al. 2023; Pavlović et al. 2022). (iii) Moreover, realistic simulation of cross-reactivity (i.e., the simulation of multiple immune signals per sequence) and chain pairing require further biological insights. (v) Finally, simulations on the repertoire level may be refined to render the simulation of public clone occurrence more native-like (Kanduri et al., n.d.), as well as by (vi) allowing for the simulation of somatic hypermutation (Yaari et al. 2013; Safra et al. 2023; Sheng et al. 2017) and (vi) associated phylogenetic lineage trees (Yermanos et al. 2017; Ralph and Matsen 2022; Hoehn, Pybus, and Kleinstein 2022; Hoehn et al. 2019; Zhang et al. 2022). There is a potential risk that an AIRR-ML model trained on simulated data may only learn the artifacts of the simulation framework, thereby impacting the applicability of ML model-related insights to real-world scenarios. This may be addressed in the future by analyzing how ML results on synthetic data transfer to experimental data as recently demonstrated by Robert et al. (Robert et al. 2022). Further work is needed on the biological understanding of immune signals for improved simulation (Kanduri et al. 2022; Widrich et al. 2020). Moreover, current speed bottleneck computations, such as those of generation probability, may be alleviated by novel encoding algorithms (Konstantinovsky and Yaari 2023)), thus improving overall simulation speed.

Future progress in AIRR-ML research is required in areas as diverse as evaluating the impact of sample size on prediction accuracy (Pavlović et al. 2021; Kanduri et al. 2022), negative dataset definition (Montemurro, Jessen, and Nielsen 2022; Deng et al. 2023; Dens et al. 2023), and unbiased estimation of prediction accuracy (Meysman et al. 2023; Moris et al. 2021b). For all of these use cases, LIgO simulations may be employed for benchmarking and developing interpretable AIRR-ML methods.

## Methods

### Simulation of background AIRs

LIgO supports two sources for background AIRs — (i) simulation of synthetic AIRs using a V(D)J recombination model, or (ii) processing experimental or any other set of AIRs provided by the user. The chosen source of AIRs will also be used during the simulation of immune-signal-specific AIRs.

To simulate naive synthetic AIRs, LIgO uses the OLGA tool (Sethna et al. 2019) according to a predefined V(D)J recombination model (OLGA model), but, in principle, any other V(D)J recombination simulator such as IGoR (Marcou, Mora, and Walczak 2018) or immuneSIM (C. R. Weber et al. 2020) can be used. Briefly, OLGA describes a V(D)J recombination using a stochastic process, whose parameters include the distribution of V genes (or alleles), a conditional probability of J genes (or alleles) given a V gene (or allele), and others. OLGA produces AIRs in the format of a V gene (or allele) name, CDR3 sequence (nucleotide and amino acid), and a J gene (or allele) name, for example, TRBV3*01 + CASSLGGVGYEQYF + TRBJ2-7*01. LIgO supports the usage of the default OLGA models with parameters estimated using IGoR (Marcou, Mora, and Walczak 2018). This includes human AIRs (IGH, IGL, IGK, TRA, TRB) and mice AIRs (TRA, TRB). Alternatively, the user may specify a custom OLGA model. For example, a user can simulate mosaic deletion patterns if they assign (conditional) probabilities of zero to several gene alleles (Gidoni et al. 2019).

The second option for the simulation of background AIRs is a user-provided set of annotated AIRs in the following formats — AIRR Community standard format (Rubelt et al. 2017), MIXCR (Bolotin et al. 2015), 10x Genomics (Zheng et al. 2017), Adaptive Biotechnologies ImmunoSEQ (Emerson et al. 2017; Nolan et al. 2020), or VDJdb formats (Bagaev et al. 2020). In this case, LIgO will iterate through every AIR once until the set of AIRs is empty. If the user-provided set of AIRs is not large enough to complete the simulation, LIgO will inform the user that a larger pool of background AIRs is needed to complete the simulation.

### Representation of motifs

LIgO allows for defining motifs in three different ways (Fig. 1C) — k-mers, gapped k-mers, and PWMs (positional weight matrices). K-mers and gapped k-mers might be more convenient for initial proof-of-concept simulations as they are more straightforward to define than PWMs.

1. A *k-mer* is defined as a nucleotide or amino acid subsequence of length k, for example, AAA. Theoretically, a k-mer can be an entire CDR3 region. LIgO can also introduce several mutations to a k-mer defined by a Hamming distance range, i.e., the number of allowed mutations from the original k-mer to the mutated k-mer. For example, a k-mer AAAA with additional hamming distance range from 1 to 2 corresponds to a set of 4-mers that contain two or three As and the other positions may be any nucleotide or amino acid.
2. A *gapped k-mer* is defined as a nucleotide or amino acid subsequence that contains k nucleotides or amino acids and gap(s). In LIgO, the gap is allowed only in one position and a gapped k-mer is parameterized with a subsequence, gap position, and gap length range. For example, a gapped 3-mer AA/A with a gap range from 1 to 3 encodes the subsequences AA-A, AA--A, and AA---A, where a gap, denoted with “-”, corresponds to one arbitrary nucleotide or amino acid. Similarly to k-mers, LIgO supports the definition of hamming distance with a gapped k-mer.
3. A *PWM of length k is* defined as a joint probability distribution of k independent discrete distributions over nucleotides or amino acids. Every PWM column describes a distribution for a given position of a motif. A k-mer is a special case of a PWM where the values in every column are zero except for one nucleotide or amino acid that is 1. A gapped k-mer is a PWM whose values in every column are either 0 or 1 in the positions where there is no gap, and uniformly distributed among nucleotides or amino acids in the gap positions.

A critical concept for LIgO simulation is defining whether a given AIR contains a motif. If a motif is a k-mer or a gapped k-mer, an AIR must include this (gapped) k-mer as a substring of the CDR3 region. A (gapped) k-mer or a PWM search in an AIR is implemented using BioNumPy (Rand et al. 2022) enabling efficient representation and operations with AIRs. Additionally, the search for (gapped) k-mers can accommodate mismatches, allowing for hamming distance consideration. If a motif is a PWM, LIgO constructs a regular expression based on the PWM and checks if an AIR matches the regular expression. For every subsequence matching the regular expression, LIgO calculates a PWM score, which equals its log-likelihood under the distribution described by PWM. Only those subsequences with a PWM score above a user-defined threshold are considered a match.

PWMs and (gapped) k-mers can describe various motifs in a sequence. However, using only one PWM, we cannot introduce dependencies between different positions in a subsequence because, by definition, the discrete distributions corresponding to every motif position are independent. Therefore several PWMs or (gapped) k-mers may have to be combined to describe an immune signal and capture the biological complexity. For example, if one were to define a motif A(A/C)(A/C) where only one C is allowed, one would have to combine two k-mers AAC and ACA.

### Representation of immune signals

Immune signals (Fig. 1C) combine motifs with positional dependencies and immune-specific information. LIgO implements two types of immune signals: (i) a set of motifs that are (optionally) restricted with the V and J gene or allele names and the position(s) in the CDR3. For example, a set of motifs {AAA, CCC, A--CC} restricted by IGHV1*01 and IGHJ4, which occur only on the IMGT positions 108 or 110 (Lefranc et al. 2003). In this case, an AIR contains an immune signal if it contains any of the motifs on a particular CDR3 position and the correct V and J gene, if the IMGT position(s) or V/J genes are defined. (ii) The second way to define an immune signal is a custom function indicating the signal’s presence in an AIR. The function takes the amino acid and the nucleotide AIR sequence, its V and J gene and returns true if the AIR contains the desired signal and false otherwise. Such custom functions allow users to define any immune signal. For example, for an antigen-specific ML classifier that takes a sequence as input and produces the signal label as output (Mason et al. 2021; Montemurro, Jessen, and Nielsen 2022; Akbar, Robert, et al. 2022), the user could write a custom function to call the classifier and define an immune signal based on the classifier’s output.

### Representation of immune events

Immune events (Fig. 1C), such as diseases, vaccinations, or allergies, elicit immune responses which change an individual’s adaptive immune receptor repertoire. AIRRs with the same immune event might share similar immune signals involved in the immune response. More formally, LigO defines an immune event as a set of immune signals and their proportion in an AIRR. For example, we can define an AIRR with T1D (type 1 diabetes) experience as containing 1% of receptors with corresponding signals. To reflect biological complexity, LIgO-simulated immune repertoires may contain multiple immune events.

For a receptor-level simulation, an AIR belongs to an immune event if it contains any of the signals for the immune event. Implementation-wise, the immune event for a single AIR is the next level of abstraction after the immune signal and is directly assigned to an AIR. It does not influence the AIR sequence in any way, but logically, it should follow the immune signals that are considered to belong to a given immune event. Since it does not influence the simulation, there might be as many immune event labels as desired. For example, we can regard an immune event T1D as a set of two immune signals {signal1 = “AAA”, signal2 = “GGG”}. In this case, an AIR (TRBV13*01+ CASSAGGGAFYGYTF + TRBJ1-2*01) will be signal2-specific because it contains “GGG”. The user may then choose to explicitly assign the “T1D” label to the receptor. From a practical standpoint, a user would define a group of receptors with a particular set of simulation parameters and labels.

### Generalization of the immune signal concept to paired-chain AIRR data

Both BCRs and TCRs are (generally) composed of two protein chains — heavy and light chains for BCRs and alpha/beta or gamma/delta chains for TCRs. Until recently, most studies focused on the heavy chain of BCRs and beta or delta chains of TCRs (Zhou and Kleinstein 2019; Georgiou et al. 2014). However, the pairing information is important for studying AIR diversity (Shcherbinin, Belousov, and Shugay 2020) and antigen binding (Rossjohn et al. 2015). Recently developed single-cell AIRR-seq technologies enable accurate identification of paired AIRs (Stubbington et al. 2017; Friedensohn, Khan, and Reddy 2017). Currently, it remains unclear whether the pairing rules that combine two chains in one receptor follow a uniform distribution or not, and until now, there are no studies that were able to infer the pairing rules (Dupic et al. 2019; Jayaram, Bhowmick, and Martin 2012; B. DeKosky 2017; B. J. DeKosky et al. 2016). Furthermore, there is little evidence of major structural constraints in chain pairing (Tanno et al. 2020).

The LIgO simulation framework is based on the simplifying assumption that an AIR is independently formed by two chains, which allows processing these two chains separately and defining an immune signal of a paired AIR as immune signals of each two receptor sequences. We define that a paired chain is part of an immune signal if and only if at least one chain contains the immune signal. A special case of a paired immune signal is a pair of two signals where one or even two AIRs are predefined (i.e., a corresponding motif is a whole receptor). In this case, all the paired receptors that are part of this paired immune signal will share a similar or same light chain. A paired immune event is defined as a pair of a specificity label and a set of paired immune signals.

Simulation of paired signals is performed similarly to the single-chain signal simulation described above. Each chain is independently simulated using either rejection sampling or signal implantation. After that, single chains are grouped into pairs per user specification.

### Immune receptor annotation with immune signals

LIgO annotates an AIR as containing an immune signal or not, via checking if any of the immune signals defined in the simulation occurs anywhere in the AIR where it’s allowed (per IMGT positions if they are specified). If one AIR contains two overlapping motifs from different signals, the AIR is labeled as specific to both signals and removed if two signals per receptor are not specified for that simulation batch. If one AIR contains two motifs from the same signal, they are not considered separately and the AIR is labeled as specific to that signal.

### Simulation of immune-signal-specific receptors using rejection sampling

Rejection sampling enables simulation of event-specific AIRs without changing their sequence. The rejection sampling procedure is performed iteratively. During each iteration, a batch of background AIRs is selected either from the background V(D)J model or experimental data. By default, the batch size is 10^4^ AIRs. If an immune signal contains signals which are restricted with a specific V or J gene and the simulation is defined by a V(D)J model, LIgO uses a skewed V(D)J model to simulate AIRs containing only the predefined V and J genes to speed up the simulation process. After that, each AIR from the batch is annotated with all immune signals. AIRs containing more than two immune signals are eliminated. AIRs containing one immune signal or a combination of two immune signals (as per user specification) are added to a pool of immune-signal-specific receptors. The process is repeated until the desired amount of receptors is not simulated for all immune signal(s) or the number of iterations exceeds the upper limit, which is 100 by default.

### Simulation of immune-signal-specific receptors using signal implantation

Signal implantation is the fastest way to simulate immune-signal-specific receptors. However, this method is limited to immune signals which are defined as either a (gapped) k-mer or PWM. This limitation arises because implantation of other signal components such as a V gene or CDR3 length, may impact an AIR sequence structure. Once the user defines the desired amount of AIRs for each immune signal, LIgO obtains this specified amount of background receptors from either a recombination model or user-defined experimental data. After that, the corresponding (gapped) k-mer or a substring generated by a PWM is implanted to each AIR, respectively. Implantation can be performed at any position of the CDR3 region in such a way that the implanted subsequence does not increase the length of a given AIR. The implantation position can be uniformly distributed across the CDR3 or may be defined by the user according to a distribution across IMGT positions. For instance, positions in the center of the CDR3 might have a high probability of implantation, while positions in the conserved regions such as the beginning and the end of the CDR3 might exhibit negligible probability.

### Generation probability evaluation

When a LIgO simulation is defined by an OLGA V(D)J recombination model (Sethna et al. 2019; Marcou, Mora, and Walczak 2018), it enables the computation of generation probability for all AIRs involved in the simulation process with respect to this specific model. The generation probability (pgen) of an AIR estimates the probability of a given OLGA model to generate a given AIR (Sethna et al. 2019). Pgen is calculated as the cumulative probability across all potential recombination scenarios capable of producing this AIR. While generation probability can provide valuable insights into AIRR analysis (Pogorelyy et al. 2019; Marcou, Mora, and Walczak 2018; Elhanati et al. 2018; Slabodkin et al. 2021), it is worth noting that pgen computation might lead to an increase in overall simulation time due to the slower nature of OLGA’s pgen calculation.

LIgO does not support pgen evaluation if a simulation is defined via user-provided experimental data. In this case, the corresponding recombination model is not specified since experimental data may be merged from multiple individuals and thus align with distinct recombination models (Slabodkin et al. 2021).

### Importance sampling

If a LIgO simulation is defined by an OLGA V(D)J recombination model, LIgO supports an importance sampling approach, which preserves the pgen distribution of a simulated AIRR close to the background pgen distribution. First, LIgO estimates the pgen distribution of background AIRs through a histogram of logarithmically transformed pgens of a large set of background AIRs. To do so, LIgO generates a batch of background AIRs, and divides these AIRs into equal-width histogram bins (in log-scale), where the number of bins must be provided by the user. Next, an acceptance probability is assigned to each bin of the histogram. The acceptance probability is proportional to how often we observe background AIRs falling into the pgen range corresponding to the histogram bin. If there are any pgen bins without observed AIRs, they are set to a predefined minimum acceptance probability of 10^-5^. All these acceptance probabilities are then normalized to sum to 1.

During the simulation of event-specific AIRs, LIgO generates an AIR either with rejection sampling or signal implantation and calculates the logarithm of its pgen. This log-pgen is then matched with the corresponding bin in the histogram and the acceptance probability associated with this bin. Subsequently, the AIR is accepted into the final dataset with a probability equal to the acceptance probability. This process is continued until the desired amount of event-specific AIRs has been simulated.

### Assessment of simulation feasibility

To evaluate the simulation feasibility for a given set of parameters, LIgO generates a batch of background AIRs, with the default batch size being 10^5^ AIRs. After that, each AIR is annotated with all immune signals participating in the simulation process and AIRs containing more than 2 immune signals are filtered out. The ratio of immune-signal specific receptors to the batch size estimates the immune signal frequency among the background receptors. If the estimated frequency of an immune signal is excessively low (less than 0.001%), then it is not recommended to use rejection sampling for simulating this immune signal. This is due to the number of background receptors that must be generated during rejection sampling to find at least one event-specific AIR is inversely proportional to the background frequency of this immune signal. Conversely, if the estimated frequency is too high (more than 10%), then such an immune signal may lack biological relevance and might cause an increased number of simulation iterations, if the user wants to add background receptors without any immune signals to the final dataset.

LIgO also estimates an immune signal co-occurrence matrix, which represents the conditional probability of observing one immune signal within an AIR given the presence of another immune signal. This conditional probability of an AIR containing immune signal 2 given the presence of immune signal 1 is calculated as a ratio of the number of AIRs within the original batch containing both immune signal 1 and immune signal 2 and the number of AIRs containing only immune signal 1. For users considering rejection sampling while ensuring each AIR holds no more than one signal, the co-occurrence matrix indicates which immune signals have high chances of co-existing in one AIR. These immune signals are recommended to be re-defined. Alternatively, users can use implantation instead of rejection sampling since it can restrict the number of immune signals to one for each AIR.

### Clonal count simulation

In a repertoire-based scenario, LIgO offers an optional simulation of clonal counts for each AIR. These counts follow the Zeta distribution, which is a discrete analog of the power law distribution. The realization of the Zeta distribution is performed using the Zipf function from the scipy.stats library (Virtanen et al. 2020), which requires two parameters — *a* (shape parameter, where a≥1) and *loc* (shift parameter). For each immune signal and the background AIRs, the user needs to define *a* and *loc* parameters. By adjusting these parameters across different immune signals, one can simulate clonal expansion of AIRs containing these specific signals (Greiff et al. 2015).

### Identification of amino acid 3-mer frequency distribution patterns in CDR3 regions of synthetic IGHs and TRBs

To demonstrate the natural distribution of 3-mers within CDR3 regions, we used LIgO to simulate a dataset of 10^6^ amino acid background TRBs using the default IGoR generation model (Marcou, Mora, and Walczak 2018; Sethna et al. 2019), where each CDR3 has a restricted length of 15 amino acids. After that, we calculated how often each 3-mer appears on each CDR3 position among simulated TRBs using immuneML (Pavlović et al. 2021). We only considered 3-mers starting at the IMGT positions 105–115, because the positions 104 and 118 are conserved and thus deliberately excluded from the analysis. We also eliminated from the analysis all 3-mers that were present in less than 10 TRBs out of the total amount of 10^6^ simulated TRBs. For each 3-mer we normalized the frequencies of this 3-mer across positions 105–115, ensuring that the total sum equals one. Normalized frequency vectors were then clustered using hierarchical clustering with complete linkage method and Euclidean distance between vectors. The hierarchical clustering dendrogram was split into 10 clusters and for each cluster, we calculated the mean frequency and 95% confidence interval among all 3-mers belonging to this cluster for all IMGT positions 105–115. Nine clusters out of ten (clusters 2-10) containing 48.69% of amino acid 3-mers in total have a peak of average frequency, i.e. average frequency > 0.4, at one of the IMGT positions 105–108 or 111–115 (Fig. 3). One cluster (cluster 1) containing 59.92% 3-mers has no average peaks.

The same analysis was also performed on 10^6^ amino acid IGHs, see Supplementary Fig. 2 for more details. Three out of ten clusters (clusters 1, 3, 4, 79.98% of 3-mers) do not contain a frequency peak on average at any IMGT position, seven out of ten clusters (clusters 2, 5–10, 19.85% of 3-mers) have an average peak at one of the IMGT positions 105–108 or 111–115.

### Signal implantation decreases the generation probability of a given receptor

We conducted a comparison of pgens before and after implantation across four different signals, where one signal is defined as a set of ten random amino acid k-mers of the same length; see Supplementary Table 2 for the list of used k-mers. We simulated four TRB datasets, each corresponding to a different k-mer size, specifically k = 2, 3, 4, and 5. For each k-mer, we simulated 100 signal-specific amino acid TRBs using implantation and the default IGoR model (Marcou, Mora, and Walczak 2018; Sethna et al. 2019), resulting in a total of 1000 TRBs for each individual dataset. The implanting positions were constrained to ensure that each implanted k-mer does not affect the first and the last two amino acids of the original CDR3 sequence. Using the default IGoR model (Marcou, Mora, and Walczak 2018; Sethna et al. 2019), we computed a pgen difference for each TRB by subtracting the log10-transformed pgen after implantation from the log10-transformed pgen before implantation. Receptors with a post-implantation pgen value of zero were excluded from the analysis, resulting in removing 0.8% of TRBs for 2-mers, 0.7% of TRBs for 3-mers, 2.4% of TRBs for 4-mers, and 3% of TRBs for 5-mers. The majority of TRBs exhibited a positive pgen difference, with percentages as follows: 95.1% for 2-mers, 96% for 3-mers, 97% for 4-mers, and 93.4% for 5-mers. The median pgen difference values were 2.46 for 2-mers, 3.04 for 3-mers, 4.14 for 4-mers, and 4.05 for 5-mers. This indicates that, on average, the pgen of a receptor after implantation may experience a reduction of between two and four orders of magnitude, depending on the length of the k-mer.

Similarly, we calculated pgen difference when immune-signal-specific AIRs are generated using signal implantation coupled with importance sampling, where the background pgen distribution was estimated using 10^4^ TRBs splitted into 30 histogram bins. In this scenario, the average pgen difference decreased compared to signal implantation performed on its own. The median pgen difference values were 1.77 for 2-mers, 2.5 for 3-mers, 3.36 for 4-mers, and 3.46 for 5-mers.

### Implantation coupled with importance sampling yields an overall similar pgen distribution to rejection sampling for signal-specific AIRs

We compared the generation probability distribution of signal-specific AIRs generated using three simulation approaches — rejection sampling (RS), implantation (I), and implantation coupled with importance sampling (I + IS). The immune signal was defined as a set of ten random k-mers of the same length, where k = 2, 3, 4, and 5; see the detailed description of the immune signals and the simulation of signal-specific AIRs using rejection sampling and signal implantation in the previous section. To simulate signal-specific AIRs using implantation coupled with importance sampling, we constructed a histogram of logarithmically transformed pgens using 30 bins and 10^4^ background TRBs simulated with the default IGoR model (Marcou, Mora, and Walczak 2018; Sethna et al. 2019). Next, we evaluated acceptance probabilities for each bin and continued the simulation until the number of accepted AIRs for each k-mer was 100. The pgen distribution of signal-specific AIRs generated using implantation coupled with importance sampling is overall closer to the rejection sampling pgen distribution, which is the ground-truth pgen distribution of signal-specific AIRs in logarithmic scale. The median of the log-pgen distribution for each method are: 3.5⋅10^-15^ (I), 8.1⋅10^-13^ (RS), 1.2⋅10^-12^ (I+IS) for 2-mers; 1.9⋅10^-15^ (I), 1.4⋅10^-13^ (RS), 1.4⋅10^-13^ (I+IS) for 3-mers; 1.4⋅10^-16^ (I), 3.3⋅10^-15^ (RS), 1.8⋅10^-14^ (I+IS) for 4-mers; and 2.7⋅10^-16^ (I), 5.6⋅10^-15^ (RS), 1.9⋅10^-14^ (I+IS) for 5-mers.

### Use case 1: Out-of-distribution receptor-level simulation using LIgO

To demonstrate how different train-test split strategies may impact the prediction accuracy and the optimal model, we simulated a dataset of 4⋅10^4^ IGH amino acid sequences in 10 replicates (4⋅10^5^ IGHs in total) using signal implantation and the default IGoR IGH V(D)J recombination model (Marcou, Mora, and Walczak 2018). Each dataset of 4⋅10^4^ IGH sequences represented a set of AIRs obtained from four different individuals, where every individual contributed 5⋅10^3^ signal-specific AIRs and 5⋅10^3^ non-specific AIRs to the dataset. The immune signal was defined as a set of four similar 4-mers — AAAA, AAGA, AACA, and AANA, and we assumed that every individual carries only one signal 4-mer out of four. Altogether, one dataset contained 10^4^ IGH sequences from person 1 (5⋅10^3^ IGH sequences with AAGA, 5⋅10^3^ IGH sequences without any of four signal 4-mers), 10^4^ IGH sequences from person 2 (5⋅10^3^ IGH sequences with AACA, 5⋅10^3^ IGHs without without any of four signal 4-mers), 10^4^ IGH sequences from person 3 (5⋅10^3^ IGH sequences with AANA, 5⋅10^3^ IGH sequences without without any of four signal 4-mers), and 10^4^ IGH sequences from person 4 (5⋅10^3^ IGH sequences with AAAA, 5⋅10^3^ IGH sequences without without any of four signal 4-mers), where non-specific IGHs did not contain any of the four signal 4-mers.

We used a logistic regression model to perform a binary receptor-based classification task, i.e., classify AIRs into a signal-specific class or non-specific class. You can find more details about our usage of logistic regression in the “Machine Learning” methods section. The logistic regression model was trained and tested using the data described above and two train-test split approaches — (i) random train-test split and (ii) leave-one individual out train-test split. For the random train-test split strategy, 4⋅10^4^ IGH sequences were randomly split into 75% train and 25% test subsets four times for the model assessment on the outer cross-validation loop, and every time within the inner cross-validation loop, threefold cross-validation was used for the model (e.g., hyperparameter) selection. For the leave-one-individual out procedure, we split 10^4^ AIRs from one individual to the test set and the rest 3⋅10^4^ AIRs to the train set for the model assessment (outer) cross-validation loop and this was repeated for every individual. In the model selection (inner) cross-validation loop we again split 10^4^ AIRs from one individual to the test set, and the rest 2⋅10^4^ AIRs to the train set and repeated this procedure for every individual.

We benchmarked the logistic regression model from the scikit-learn package (Pedregosa et al. 2012) using immuneML (Pavlović et al. 2021). We encoded the IGH sequences using 4-mers encoding, where each vector of 4-mer frequencies was normalized by scaling its variance to 1, and mean to zero across different IGH sequences. We used two train-test strategies and performed hyperparameter optimization for regularization parameters (L1 and L2), and the regularization constant (0.01, 0.05, 0.1, 1), where the optimized performance metric during model training was balanced accuracy. We repeated the logistic regression training 10 times for every dataset replica.

For both train-test split strategies, the optimal balanced accuracy was achieved on the L1 regularization and the regularization constant 0.1. However, the model trained using the random train-test split strategy, achieved a mean balanced accuracy of 0.98, and the model trained using the leave-one-individual out achieved a mean balanced 0.67. We examined the ability of both logistic regression models to recover the ground-truth implanted 4-mers. We considered the largest absolute coefficients of the optimal logistic regression models, and the random LR models recovered all four signal 4-mers, while the leave-one-individual out models only recovered three signal 4-mers (AAGA, AANA, AACA) out of four.

### Use case 2: Limitations of conventional encoding schemes for repertoire-level binary classification when immune signals co-occur within the same AIR

We simulated 200 TRB repertoires (100 repertoires in each class and 10^5^ amino acid AIRs in each repertoire) in six replicates (three replicates for two signal frequencies 0.01% and 0.1%). We defined an immune signal as a combination of two 3-mers GDT and SGL. Both GDT and SGL occur in the naive AIRs simulated via the default IGoR TRB model (Marcou, Mora, and Walczak 2018) at approximately 0.01% frequency and co-occur in the same AIR at approximately 0.00001% frequency. Using rejection sampling, we simulated positive class AIRRs containing 0.01% or 0.1% of signal-specific receptors (containing both GDT and SGL), and the negative class containing 0.001% and 0.01% signal-specific receptors, respectively. Additionally, AIRRs from the positive and the negative classes had similar frequency of each GDT and SGL separately, which made the classes indistinguishable for the classical k-mer encoding.

Using the simulated data described above, we compared two multiple instance learning (MIL) approaches — logistic regression and a custom approach that takes into account signal co-occurrence. The first ML method has been described extensively elsewhere (Kanduri et al. 2022). Briefly, an L1-regularized logistic regression model trained on k-mer abundance-encoded data (k=3 here) was used, whose hyperparameters are optimized through nested cross-validation. Note that this method treats each k-mer in isolation and does not consider combinations. The second ML method (custom MIL) first classifies each receptor (instance) as whether immune state-associated or not and then sums up the number of positive instances per repertoire. The number of positive instances per repertoire is used as the single feature for a logistic regression. The identification of positive instances relied on the presence of pairs of 3-mers (combinations) that are together over-represented in positive class labeled receptors relative to the negative class labeled receptors as assessed through a Fisher’s exact test. For each 3-mer pair, the contingency table for Fisher’s exact test is composed of (a) the number of receptors of positive AIRRs that contain the 3-mer pair, (b) the number of receptors of positive AIRRs that do not contain the 3-mer pair, (c) the number of receptors of negative AIRRs that contain the 3-mer pair, and (d) the number of receptors of negative AIRRs that do not contain the 3-mer pair.

### Machine Learning

We utilized Logistic Regression for binary classification of signal-specific repertoires and receptors, which was similar to our previous studies (Kanduri et al. 2021; Pavlović et al. 2021). Briefly, we represented AIRs using k-mer encoding and with further standardization by subtracting the mean and dividing by the standard deviation. After that, we optimized LR hyperparameters, such as L1 and L2 regularization and regularization strength constant, using nested cross-validation. We evaluated performance of ML models using three metrics — precision 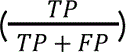, recall 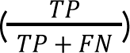, and balanced accuracy 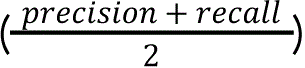, where *TP* is true positive, *FP* is false positive and *FN* is false negative. Model training, selection, and evaluation was performed using the immuneML platform (Pavlović et al. 2021) (version 2.2.3). See the Use cases methods sections for a more comprehensive description of classifiers used in every use case.

## Code availability

LIgO is freely available as a Python package from PyPI (https://pypi.org/project/ligo/), on GitHub (https://github.com/uio-bmi/ligo), and as a Docker image (https://hub.docker.com/r/milenapavlovic/ligo). The code to reproduce the manuscript figures is provided at https://github.com/mchernigovskaya/ligo-manuscript-figures.

## Declaration of interests

V.G. declares advisory board positions in aiNET GmbH, Enpicom B.V, Specifica Inc, Adaptyv Biosystems, EVQLV, Omniscope, Diagonal Therapeutics, and Absci. V.G. is a consultant for Roche/Genentech, immunai, Proteinea, and LabGenius. P.M. holds shares in ImmuneWatch BV. The remaining authors declare no competing interests.

## Funding

We acknowledge generous support by The Leona M. and Harry B. Helmsley Charitable Trust (#2019PG-T1D011, to VG), UiO World-Leading Research Community (to VG), UiO:LifeScience Convergence Environment Immunolingo (to VG, GKS, and IHH), EU Horizon 2020 iReceptorplus (#825821) (to VG), Research Council of Norway projects (#300740, 331890 to VG), a Research Council of Norway IKTPLUSS project (#311341, to VG and GKS), a Norwegian Cancer Society Grant (#215817, to VG), and Stiftelsen Kristian Gerhard Jebsen (K.G. Jebsen Coeliac Disease Research Centre) (to GKS), UiO:LifeScience Convergence Environment RealArt (to GKS, CK). This project has received funding from the Innovative Medicines Initiative 2 Joint Undertaking under grant agreement No 101007799 (Inno4Vac). This Joint Undertaking receives support from the European Union’s Horizon 2020 research and innovation programme and EFPIA (to VG). SG was supported by the Research Foundation Flanders (FWO) through an SB fellowship (1S48819N) and a travel grant for a research stay (V403821N). This work was carried out on immunohub e-Infrastructure funded by the University of Oslo and jointly operated by GreiffLab and SandveLab (the authors) in close collaboration with the University Center for Information Technology, University of Oslo, IT-Department (USIT).

## Supplementary materials

**Supplementary Table 1:** Comparison of LIgO with other AIRR simulation tools — IGoR/OLGA, immuneSIM, and simAIRR.

|  | LlgO | IGoR/OLGA | immuneSIM | simAIRR |
| --- | --- | --- | --- | --- |
| General software description |  |  |  |  |
| Reference | This work | (51, 68) | (52) | (62) |
| Purpose | Simulation of AIRR with complex immune signals for ML benchmarking (both receptor and repertoire-based) | Simulation of a set of AIRs based on a V(D)J recombination model | Simulation a set of AIRs based on a V(D)J recombination model; simulating signals (gapped k-mer only) in individual AIRs | Simulation of antigen-experienced -like AIRRs by mimicking the relation between pgen and public AIR occurrence in real-world datasets |
| Software | Python package | C++, Python package | R package | Python package |
| Simulation |  |  |  |  |
| User assistance | Yes | No | No | Yes |
| Support of both receptor and repertoire-level simulation | Receptor and repertoire level simulation | Receptor level only | Receptor level only, but supports clone counts | Repertoire level only |
| Simulation of single-chain and paired chain data | Yes | No | Yes | No |
| Immune signal complexity | Any signal complexity including (gapped) k-mers, PWMs, and user-defined signals | No | Set of k-mers | Any user-defined signals supplied in the form of full-length AIRs; strictly assumes that signal sequences are public |
| Simulation approach | Rejection sampling and signal implantation | No | Signal implantation | Implantation of full-length CDR3 amino acid sequences + V and J genes of AIRs into repertoire(s) |
| Simulates clonal expansion | Yes, somewhat (each signal and background set of AIRs may have its own clone count distribution) | No | Yes, somewhat (AIR clone count distribution may be simulated) | No |
| Methods for preserving biological nativeness of simulated data |  |  |  |  |
| Simulation based on experimental AIRR data | Yes | No, but a custom VDJ model can be used | No, but a custom VDJ model can be used | Yes |
| Preserves generation probability distribution | Yes, using importance sampling | Not applicable, the tools do not support repertoire-level simulation | No | No |
| Realistic frequencies of public AIRs in repertoire-level data when needed | No, but can be used in combination with simAIRR (see column 4 of this table) | Not applicable | Not applicable | Yes |
| Output |  |  |  |  |
| Reporting of detailed information about immune signals and the simulation | Yes | No | Yes | No |
| AIRR-compliant output format | Yes | Yes, using pygor (https://github.com/statbiophys/pygor3) | Yes | No, only OLGA format |

**Supplementary Table 2.** (relates to Fig. 3) List of random 2-mers, 3-mers, 4-mers, and 5-mers used for LIgO simulations.

| 2-mer | 3-mer | 4-mer | 5-mer |
| --- | --- | --- | --- |
| FF | KTC | NDMH | PVWGS |
| TQ | VED | VHWR | LINHT |
| CW | FFC | HDPR | RTNHL |
| LT | FPY | NSNE | QHTSV |
| DQ | CTR | YLYW | WGSDT |
| KT | VMV | CHPQ | TKDKF |
| WE | GHT | CCYN | FWMPR |
| PM | KKY | VFDP | PYLCT |
| TC | NAM | CTQA | TRYHY |
| FG | MQP | VIYM | GRLGL |

**Supplementary Figure 1.**
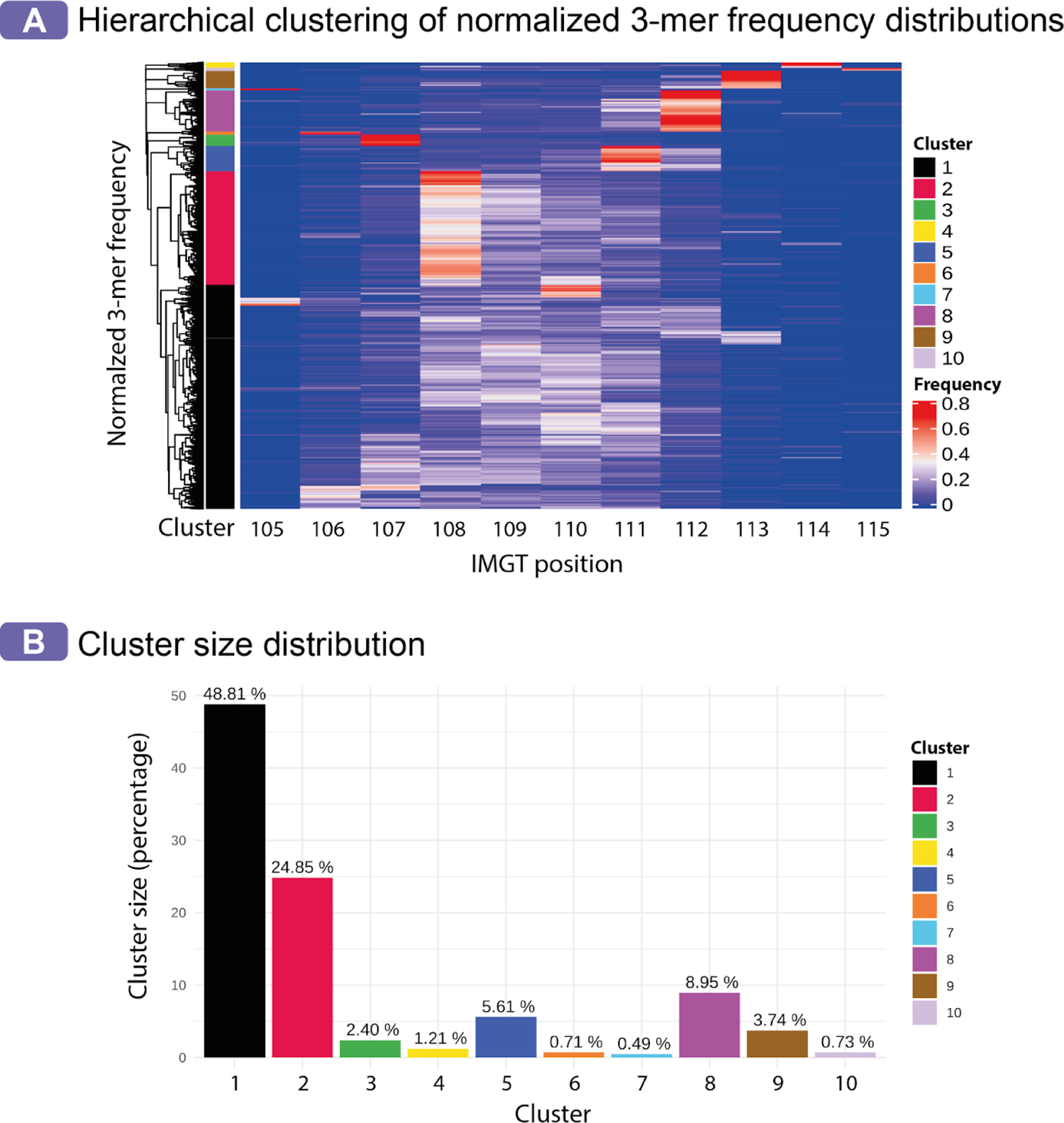
(relates to Fig. 3) Analysis of 3-mer occurrence frequency distribution clusters based on 10^6^ synthetic TRB CDR3 amino acid sequences of length 15aa. (A) Hierarchical clustering of 3-mer frequencies, where each row of the heatmap corresponds to the frequency of one 3-mer, and each column corresponds to one IMGT position. Each row (3-mer) of the heatmap is normalized such that the sum of frequencies of this 3-mer on all IMGT positions (105–115) equates to one. (B) Out of 8000 of all possible amino acid 3-mers, 48.81% of 3-mers (cluster 1) do not contain a frequency peak on average at any IMGT position, 48.69% of 3-mers (clusters 2-10) contain average frequency peak, i.e. average frequency > 0.4 at one of the IMGT positions 105–108 or 111–115, and 2.5% of 3-mers were eliminated from the analysis because they were found less than ten times in 10^6^ simulated TRB sequences.

**Supplementary Figure 2.**
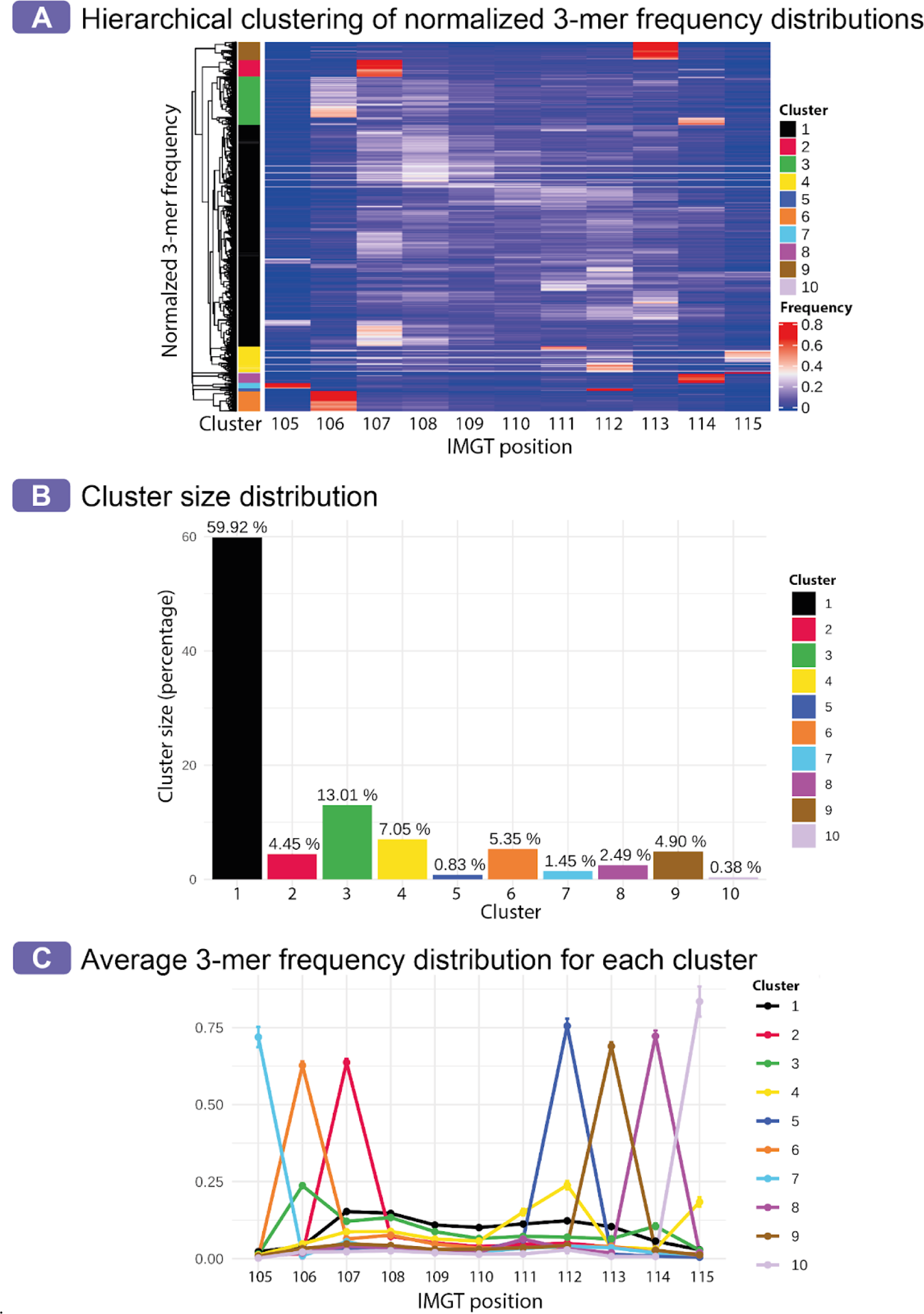
(relates to Fig. 3) Analysis of 3-mer occurrence frequency distribution clusters based on 10^6^ synthetic IGH CDR3 amino acid sequences of length 15aa. (A) Hierarchical clustering of 3-mer frequencies, where each row of the heatmap corresponds to the frequency of one 3-mer, and each column corresponds to one IMGT position. Each row (3-mer) of the heatmap is normalized such that the sum of frequencies of this 3-mer on all IMGT positions (105–115) equates to one. (B) Out of 8000 of all possible amino acid 3-mers, 79.98% of 3-mers (clusters 1, 3, 4) do not contain a frequency peak on average at any IMGT position, 19.85% of 3-mers (clusters 2, 5–10) contain an average frequency peak, i.e. average frequency > 0.4 at one of the IMGT positions 105–108 or 111–115, and 0.17% of 3-mers were eliminated from the analysis because they were found less than ten times in 10^6^ simulated TRB sequences. (C) We calculated the average frequency of all 3-mers at every IMGT position (105–115) within each cluster. One line represents one cluster, the points correspond to mean values, and the error bars correspond to the 95% confidence interval.

**Supplementary Figure 3:**
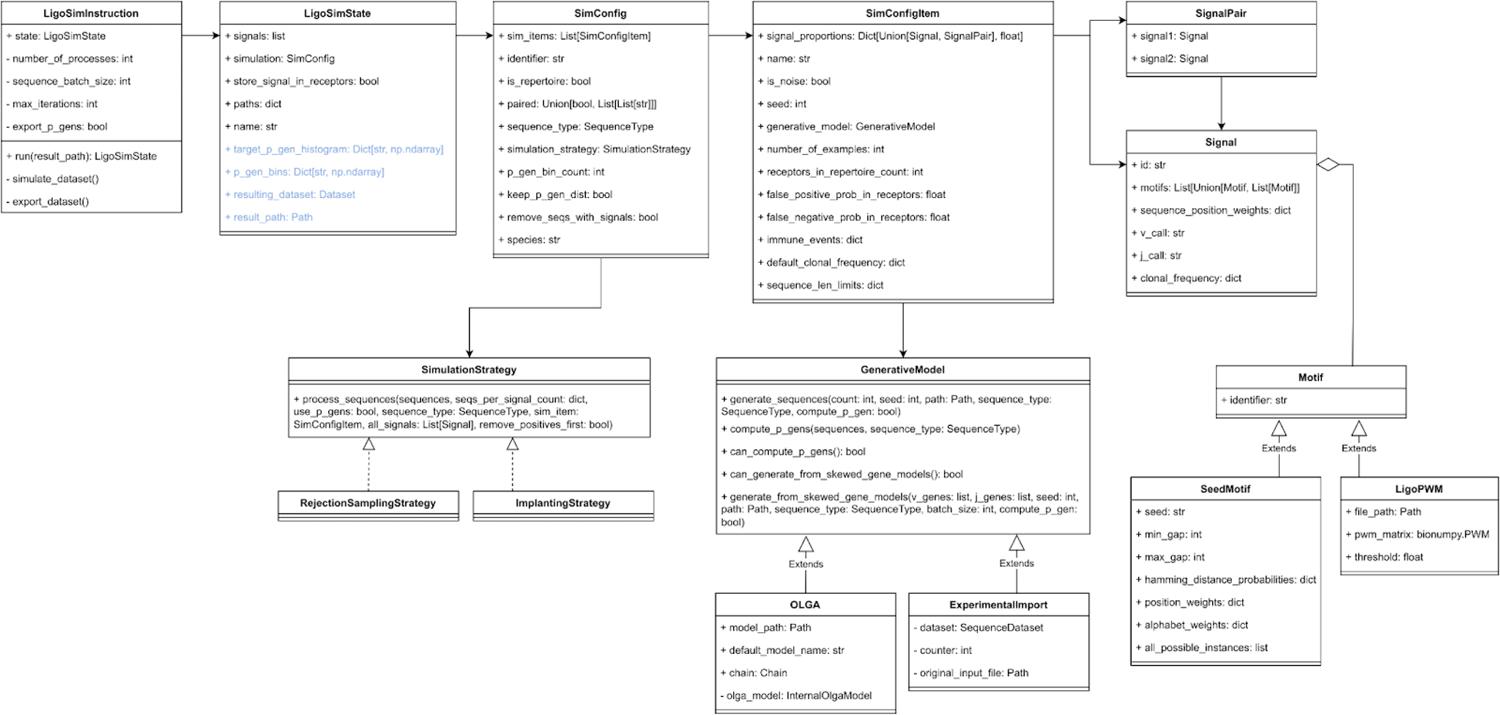
Ligo UML class diagram. The attributes in blue are set automatically at runtime, the rest are provided by the user in the YAML specification or read from default values during parsing. The LIgO data/simulation model closely follows the YAML specification.

## Notes

### Summary of Updates

References updated.

